# Common gardens in teosintes reveal the establishment of a syndrome of adaptation to altitude

**DOI:** 10.1101/563585

**Authors:** Margaux-Alison Fustier, Natalia E. Martínez-Ainsworth, Jonás A. Aguirre-Liguori, Anthony Venon, Hélène Corti, Agnès Rousselet, Fabrice Dumas, Hannes Dittberner, María G. Camarena, Daniel Grimanelli, Otso Ovaskainen, Matthieu Falque, Laurence Moreau, Juliette de Meaux, Salvador Montes-Hernández, Luis E. Eguiarte, Yves Vigouroux, Domenica Manicacci, Maud I. Tenaillon

## Abstract

In plants, local adaptation across species range is frequent. Yet, much has to be discovered on its environmental drivers, the underlying functional traits and their molecular determinants. Genome scans are popular to uncover outlier loci potentially involved in the genetic architecture of local adaptation, however links between outliers and phenotypic variation are rarely addressed. Here we focused on adaptation of teosinte populations along two elevation gradients in Mexico that display continuous environmental changes at a short geographical scale. We used two common gardens, and phenotyped 18 traits in 1664 plants from 11 populations of annual teosintes. In parallel, we genotyped these plants for 38 microsatellite markers as well as for 171 outlier single nucleotide polymorphisms (SNPs) that displayed excess of allele differentiation between pairs of lowland and highland populations and/or correlation with environmental variables. Our results revealed that phenotypic differentiation at 10 out of the 18 traits was driven by local selection. Trait covariation along the elevation gradient indicated that adaptation to altitude results from the assembly of multiple co-adapted traits into a complex syndrome: as elevation increases, plants flower earlier, produce less tillers, display lower stomata density and carry larger, longer and heavier grains. The proportion of outlier SNPs associating with phenotypic variation, however, largely depended on whether we considered a neutral structure with 5 genetic groups (73.7%) or 11 populations (13.5%), indicating that population stratification greatly affected our results. Finally, chromosomal inversions were enriched for both SNPs whose allele frequencies shifted along elevation as well as phenotypically-associated SNPs. Altogether, our results are consistent with the establishment of an altitudinal syndrome promoted by local selective forces in teosinte populations in spite of detectable gene flow. Because elevation mimics climate change through space, SNPs that we found underlying phenotypic variation at adaptive traits may be relevant for future maize breeding.

**Author summary:** Across their native range species encounter a diversity of habitats promoting local adaptation of geographically distributed populations. While local adaptation is widespread, much has yet to be discovered about the conditions of its emergence, the targeted traits, their molecular determinants and the underlying ecological drivers. Here we employed a reverse ecology approach, combining phenotypes and genotypes, to mine the determinants of local adaptation of teosinte populations distributed along two steep altitudinal gradients in Mexico. Evaluation of 11 populations in two common gardens located at mid-elevation pointed to adaptation via an altitudinal multivariate syndrome, in spite of gene flow. We scanned genomes to identify loci with allele frequency shifts along elevation, a subset of which associated to trait variation. Because elevation mimics climate change through space, these polymorphisms may be relevant for future maize breeding.

## Introduction

Local adaptation is key for the preservation of ecologically useful genetic variation [1]. The conditions for its emergence and maintenance have been the focus of a long-standing debate nourished by ample theoretical work [2–9]. On the one hand, spatially-varying selection promotes the evolution of local adaptation, provided that there is genetic diversity underlying the variance of fitness-related traits [10]. On the other hand, opposing forces such as neutral genetic drift, temporal fluctuations of natural selection, recurrent introduction of maladaptive alleles via migration and homogenizing gene flow may hamper local adaptation (reviewed in [11]). Meta-analyzes indicate that local adaptation is pervasive in plants, with evidence of native-site fitness advantage in reciprocal transplants detected in 45% to 71% of the cases [12, 13].

While local adaptation is widespread, much has yet to be discovered about the traits affected by spatially-varying selection, their molecular determinants and the underlying ecological drivers [14]. Local adaptation is predicted to increase with phenotypic, genotypic and environmental divergence among populations [6, 15, 16]. Comparisons of the quantitative genetic divergence of a trait (*Q_ST_*) with the neutral genetic differentiation (*F_ST_*) can provide hints on whether trait divergence is driven by spatially-divergent selection [17–20]. Striking examples of divergent selection include developmental rate in the common toad [21], drought and frost tolerance in alpine populations of the European silver fir [22], and traits related to plant phenology, size and floral display among populations of *Helianthus* species [23, 24]. These studies have reported covariation of physiological, morphological and/or life-history traits across environmental gradients which collectively define adaptive syndromes. Such syndromes may result from several non-exclusive mechanisms: plastic responses, pleiotropy, non-adaptive genetic correlations among traits (constraints), and joint selection of traits encoded by different sets of genes resulting in adaptive correlations. In some cases, the latter mechanism may involve selection and rapid spread of chromosomal inversions that happen to capture multiple locally favored alleles [25] as exemplified in the monkey flower, *Mimulus guttatus* [26]. While distinction between these mechanisms is key to decipher the evolvability of traits, empirical data on the genetic bases of correlated traits are currently lacking [27].

The genes mediating local adaptation are usually revealed by genomic regions harboring population-specific signatures of selection. These signatures include alleles displaying greater-than-expected differentiation among populations [28] and can be identified through *F_ST_*-scans [29–35]. However, *F_ST_*-scans and its derivative methods [28] suffer from a number of limitations, among them a high number of false positives (reviewed in [36, 37]) and the lack of power to detect true positives [38]. Despite these caveats, *F_ST_*-outlier approaches have helped in the discovery of emblematic adaptive alleles such as those segregating at the *EPAS1 locus* in Tibetan human populations adapted to high altitude [39]. An alternative to detect locally adaptive loci is to test for genotype-environment correlations [35, 40–45]. Correlation-based methods can be more powerful than differentiation-based methods [46], but spatial autocorrelation of population structure and environmental variables can lead to spurious signatures of selection [47].

Ultimately, to identify the outlier loci that have truly contributed to improve local fitness, a link between outliers and phenotypic variation needs to be established. The most common approach is to undertake association mapping. However, recent literature in humans has questioned our ability to control for sample stratification in such approach [48]. Detecting polymorphisms responsible for trait variation is particularly challenging when trait variation and demographic history follow parallel environmental (geographic) clines. Plants however benefit from the possibility of conducting replicated phenotypic measurements in common gardens, where environmental variation is controlled. Hence association mapping has been successfully employed in the model plant species *Arabidopsis thaliana*, where broadly distributed ecotypes evaluated in replicated common gardens have shown that fitness-associated alleles display geographic and climatic patterns indicative of selection [49]. Furthermore, the relative fitness of *A. thaliana* ecotypes in a given environment could be predicted from climate-associated SNPs [50]. While climatic selection over broad latitudinal scales produces genomic and phenotypic patterns of local adaptation in the selfer plant *A. thaliana*, whether similar patterns exist at shorter spatial scale in outcrossing species remains to be elucidated.

We focused here on a well-established outcrossing plant system, the teosintes, to investigate the relationship of molecular, environmental, and phenotypic variation in populations sampled across two elevation gradients in Mexico. The gradients covered a relatively short yet climatically diverse, spatial scale. They encompassed populations of two teosinte subspecies that are the closest wild relatives of maize, *Zea mays* ssp. *parviglumis* (hereafter *parviglumis*) and *Z. mays* ssp. *mexicana* (hereafter *mexicana*). The two subspecies display large effective population sizes [51], and span a diversity of climatic conditions, from warm and mesic conditions below 1800 m for *parviglumis*, to drier and cooler conditions up to 3000 m for *mexicana* [52]. Previous studies have discovered potential determinants of local adaptation in these systems.

At a genome-wide scale, decrease in genome size correlates with increasing altitude, which likely results from the action of natural selection on life cycle duration [53, 54].

More modest structural changes include megabase-scale inversions that harbor clusters of SNPs whose frequencies are associated with environmental variation [55, 56]. Also, differentiation- and correlation-based genome scans in teosinte populations have succeeded in finding outlier SNPs potentially involved in local adaptation [57, 58]. But a link with phenotypic variation has yet to be established.

In this paper, we genotyped a subset of these outlier SNPs on a broad sample of 28 teosinte populations, for which a set of neutral SNPs was also available; as well as on an association panel encompassing 11 populations. We set up common gardens in two locations to evaluate the association panel for 18 phenotypic traits over two consecutive years. Individuals from this association panel were also genotyped at 38 microsatellite markers to enable associating genotypic to phenotypic variation while controlling for sample structure and kinship among individuals. We addressed three main questions: What is the extent of phenotypic variation within and among populations? Can we define a set of locally-selected traits that constitute a syndrome of adaptation to altitude? What are the genetic bases of such syndrome? We further discuss the challenges of detecting phenotypically-associated SNPs when trait and genetic differentiation parallel environmental clines.

## Results

### Trait-by-trait analysis of phenotypic variation within and among populations

In order to investigate phenotypic variation, we set up two common garden experiments located in Mexico to evaluate individuals from 11 teosinte populations (Fig 1). The two experimental fields were chosen because they were located at intermediate altitudes (S1 Fig). Although natural teosinte populations are not typically encountered around these locations [52], we verified that environmental conditions were compatible with both subspecies (S2 Fig).The 11 populations were sampled among 37 populations (S1 Table) distributed along two altitudinal gradients that range from 504 to 2176 m in altitude over ∼460 kms for gradient *a*, and from 342 to 2581m in altitude over ∼350 kms for gradient *b* (S1 Fig). Lowland populations of the subspecies *parviglumis* (n=8) and highland populations of the subspecies *mexicana* (n=3) were climatically contrasted as can be appreciated in the Principal Component Analysis (PCA) computed on 19 environmental variables (S2 Fig). The corresponding set of individuals grown from seeds sampled from the 11 populations formed the association panel.

**Figure 1.**
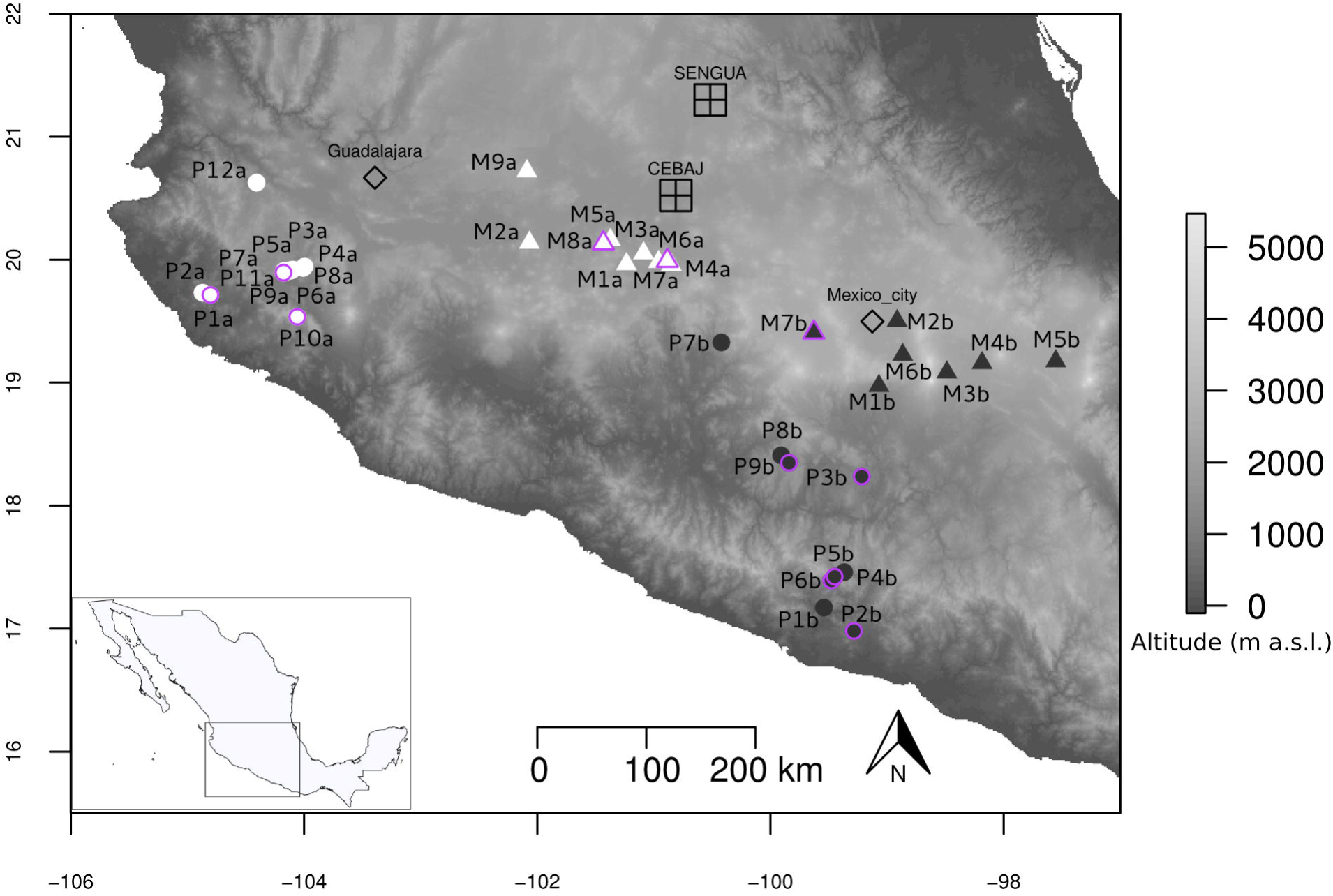
Geographical location of sampled populations and experimental fields. The entire set of 37 Mexican teosinte populations is shown with *parviglumis* (circles) and *mexicana* (triangles) populations sampled along gradient *a* (white) and gradient *b* (black). The 11 populations indicated with a purple outline constituted the association panel. This panel was evaluated in a four-block design over two years in two experimental fields located at mid-elevation, SENGUA and CEBAJ. Two major cities (Mexico City and Guadalajara) are also indicated. Topographic surfaces have been obtained from International Centre for Tropical Agriculture (Jarvis A., H.I. Reuter, A. Nelson, E. Guevara, 2008, Hole-filled seamless SRTM data V4, International Centre for Tropical Agriculture (CIAT), available from http://srtm.csi.cgiar.org).

We gathered phenotypic data during two consecutive years (2013 and 2014). We targeted 18 phenotypic traits that included six traits related to plant architecture, three traits related to leaves, three traits related to reproduction, five traits related to grains, and one trait related to stomata (S2 Table). Each of the four experimental assays (year-field combinations) encompassed four blocks. In each block, we evaluated one offspring (half-sibs) of ∼15 mother plants from each of the 11 teosinte populations using a semi-randomized design. After filtering for missing data, the association panel included 1664 teosinte individuals. We found significant effects of Field, Year and/or their interaction for most traits, and a highly significant Population effect for all of them (model 1, S3 Table).

We investigated the influence of altitude on each trait independently. All traits, except for the number of nodes with ears (NoE), exhibited a significant effect of altitude (S3 Table, model 4). Note that after accounting for elevation, the population effect remained significant for all traits, suggesting that factors other than altitude contributed to shape phenotypic variation among populations. Traits related to flowering time and tillering displayed a continuous decrease with elevation, and traits related to grain size increased with elevation (Fig 2 & S3 Fig). Stomata density also diminished with altitude (Fig 2). In contrast, plant height, height of the highest ear, number of nodes with ear in the main tiller displayed maximum values at intermediate altitudes (highland *parviglumis* and lowland *mexicana*) (S3 Fig).

**Figure 2:**
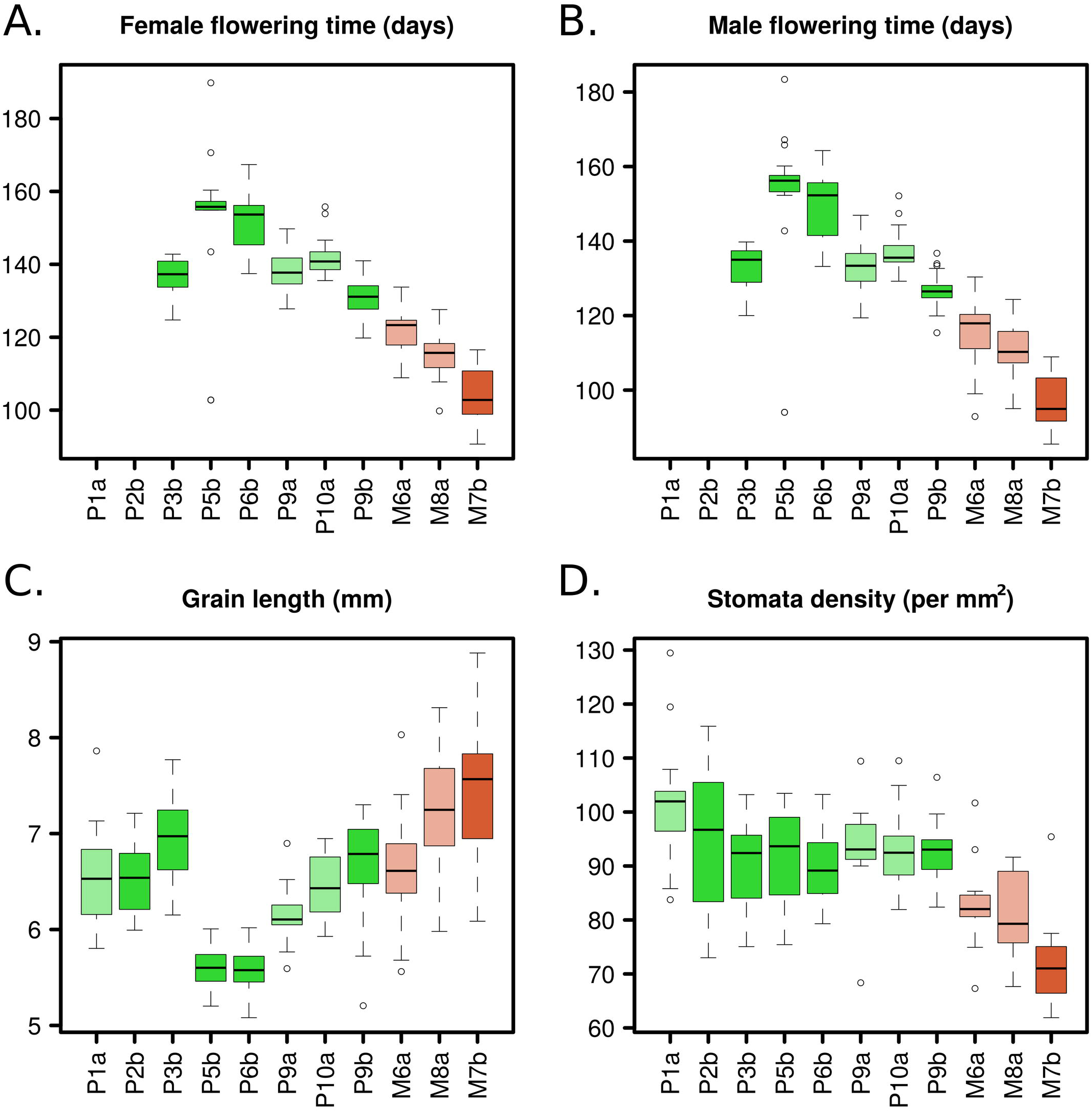
Population-level box-plots of adjusted means for four traits. Traits are female flowering time (A), male flowering time (B), grain length (C) and stomata density (D). Populations are ranked by altitude. *Parviglumis* populations are shown in green and *mexicana* in red, lighter colors are used for gradient ‘a’ and darker colors for gradient ‘b’. In the case of male and female flowering time, we report data for 9 out of 11 populations because most individuals from the two lowland populations (P1a and P2b) did not flower in our common gardens. Covariation with elevation was significant for the four traits. Corrections for the experimental setting are detailed in the Material and Methods section (Model 2).

We estimated narrow-sense heritabilites (additive genotypic effect) per population for all traits using a mixed animal model. Average per-trait heritability ranged from 0.150 for tassel branching to 0.664 for female flowering time, albeit with large standard errors (S2 Table). We obtained higher heritability for grain related traits when mother plant measurements were included in the model with 0.631 (*sd*= 0.246), 0.511 (*sd*= 0.043) and 0.274 (*sd*= 0.160) for grain length, weight and width, respectively, suggesting that heritability was under-estimated for other traits where mother plant values were not available.

### Multivariate analysis of phenotypic variation and correlation between traits

Principal component analysis including all phenotypic measurements highlighted that 21.26% of the phenotypic variation scaled along PC1 (Fig 3A), a PC axis that is strongly collinear with altitude (Fig 3B). Although populations partly overlapped along PC1, we observed a consistent tendency for population phenotypic differentiation along altitude irrespective of the gradient (Fig 3C). Traits that correlated the most to PC1 were related to grain characteristics, tillering, flowering and to a lesser extent to stomata density (Fig 3B). PC2 correlated with traits exhibiting a trend toward increase-with-elevation within *parviglumis*, but decrease-with-elevation within *mexicana* (Fig 3D). Those traits were mainly related to vegetative growth (Fig 3B). Together, both axes explained 37.4% of the phenotypic variation.

**Figure 3:**
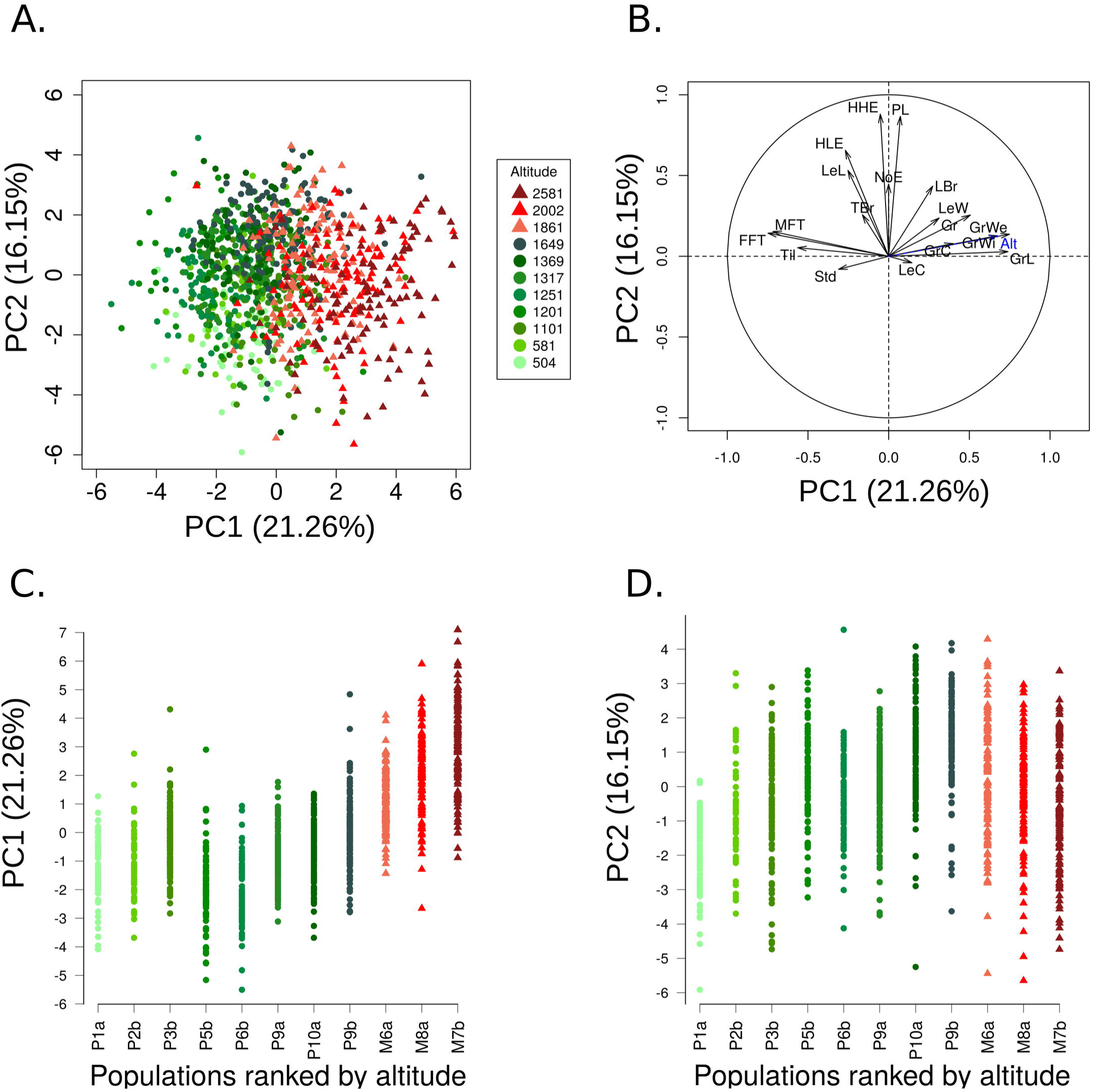
Principal Component Analysis on phenotypic values corrected for the experimental setting. Individuals factor map (A) and corresponding correlation circle (B) on the first two principal components with altitude (Alt) added as a supplementary variable (in blue). Individual phenotypic values on PC1 (C) and PC2 (D) are plotted against population ranked by altitude and color-coded following A. For populations from the two subspecies, *parviglumis* (circles) and *mexicana* (triangles), color intensity indicates ascending elevation in green for *parviglumis* and red for *mexicana*. Corrections for experimental setting are detailed in the Material and Methods (Model 3).

We assessed more formally pairwise-correlations between traits after correcting for the experimental design and population structure (*K*=5). We found 82 (54%) significant correlations among 153 tested pairs of traits. The following pairs of traits had the strongest positive correlations: male and female flowering time, plant height and height of the highest ear, height of the highest and lowest ear, grain length with grain weight and width (S4 Fig). The correlation between flowering time (female or male) with grain weight and length were among the strongest negative correlations (S4 Fig).

### Neutral structuring of the association panel

We characterized the genetic structure of the association panel using SSRs. The highest likelihood from Bayesian classification was obtained at *K*=2 and *K*=5 clusters (S5 Fig). At *K*=2, the clustering separated the lowland of gradient *a* from the rest of the populations. From *K*=3 to *K*=5, a clear separation between the eight *parviglumis* and the three *mexicana* populations emerged. Increasing *K* values finally split the association panel into the 11 populations it encompassed (S6 Fig). The *K*=5 structure reflected both altitude (lowland *parviglumis* versus highland *mexicana*) and gradients *a* and *b* (Fig 4A & B). TreeMix analysis for a subset of 10 of these populations confirmed those results with an early split separating the lowlands from gradient *a* (cf. *K*=2, S6 Fig) followed by the separation of the three *mexicana* from the remaining populations (Fig 4C). TreeMix results further supported three migration edges, a model that explained 98.75% of the variance and represented a significant improvement over a model without admixture (95.7%, Figure S7). This admixture model was consistent with gene flow between distant lowland *parviglumis* populations from gradient *a* and *b*, as well as between *parviglumis* and *mexicana* populations (Fig 4C). Likewise, Structure analysis also suggested admixture among some of the lowland populations, and to a lesser extent between the two subspecies (Fig 4B).

**Figure 4:**
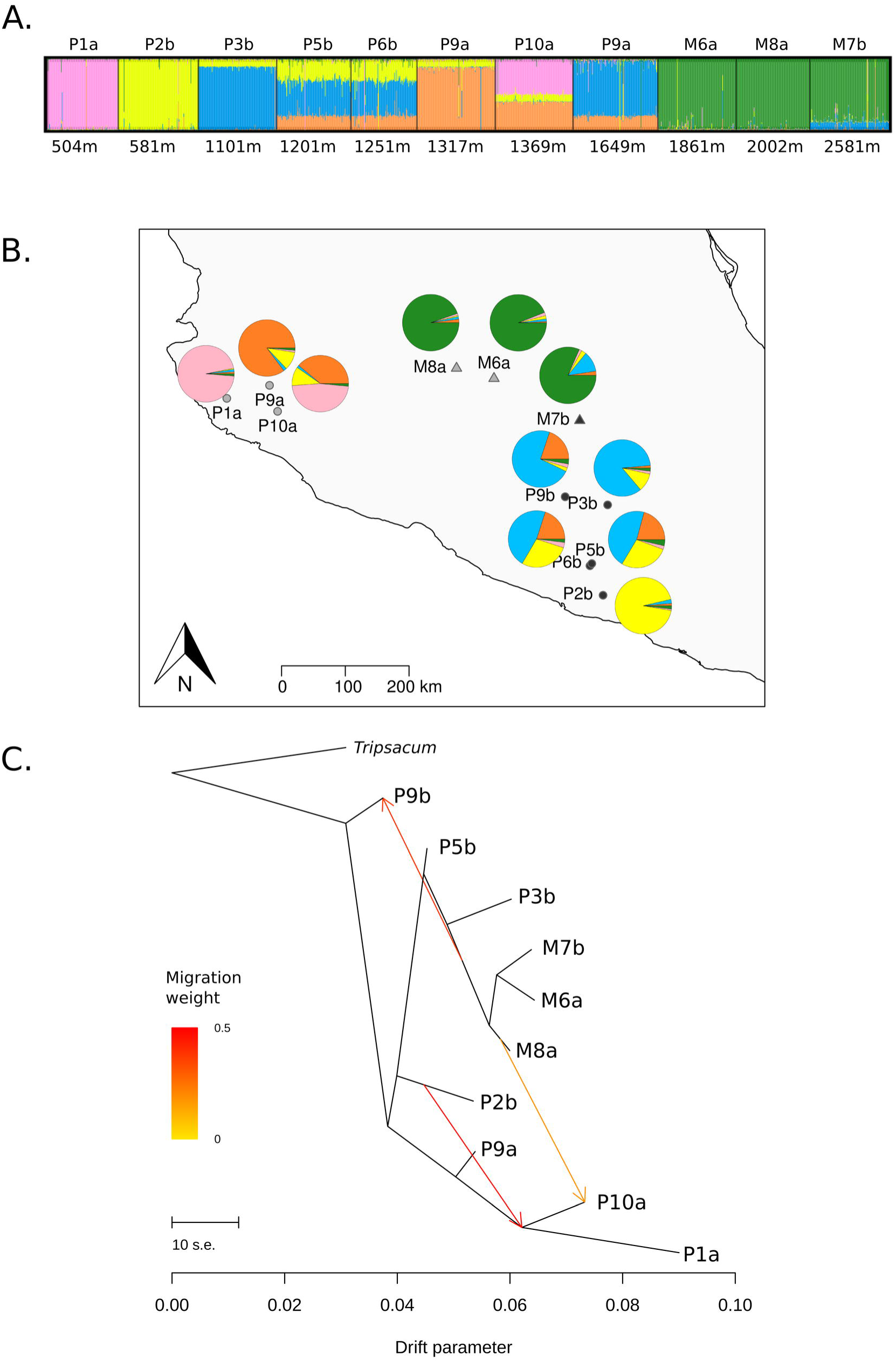
Genetic clustering, historical splits and admixture among populations of the association panel. Genetic clustering visualization based on 38 SSRs is shown for *K=5* (A). Colors represent the *K* clusters. Individuals (vertical lines) are partitioned into colored segments whose length represents the membership proportions to the *K* clusters. Populations (named after the subspecies M: *mexicana,* P: *parviglumis* and gradient ‘a’ or ‘b’) are ranked by altitude indicated in meters above sea level. The corresponding geographic distribution of populations along with their average membership probabilities are plotted (B). Historical splits and admixture between populations were inferred from neutral SNP data for a subset of 10 populations of the association panel (C). Admixtures are colored according to their weight.

### Identification of traits evolving under spatially-varying selection

We estimated the posterior mean (and 95% credibility interval) of genetic differentiation (*F_ST_*) among the 11 populations of the association panel using DRIFTSEL. Considering 1125 plants for which we had both individual phenotypes and individual genotypes for 38 SSRs (S4 Table), we estimated the mean *F_ST_* to 0.22 (0.21-0.23). Note that we found a similar estimate on a subset of 10 of these populations using 1000 neutral SNPs (*F_ST_*(CI)=0.26 (0.25-0.27)). To identify traits whose variation among populations was driven primarily by local selection, we employed the Bayesian method implemented in DRIFTSEL, that infers additive genetic values of traits from a model of population divergence under drift [59]. Selection was inferred when observed phenotypic differentiation exceeded neutral expectations for phenotypic differentiation under random genetic drift. Single-trait analyses revealed evidence for spatially-varying selection at 12 traits, with high consistency between SSRs and neutral SNPs (Table 1). Another method that contrasted genetic and phenotypic differentiation (*Q_ST_-F_ST_*) uncovered a large overlap with nine out of the 12 traits significantly deviating from the neutral model (Table 1) and one of the remaining ones displaying borderline significance (Plant height=PL, S8 Fig). Together, these two methods indicated that phenotypic divergence among populations was driven by local selective forces.

**Table 1.**
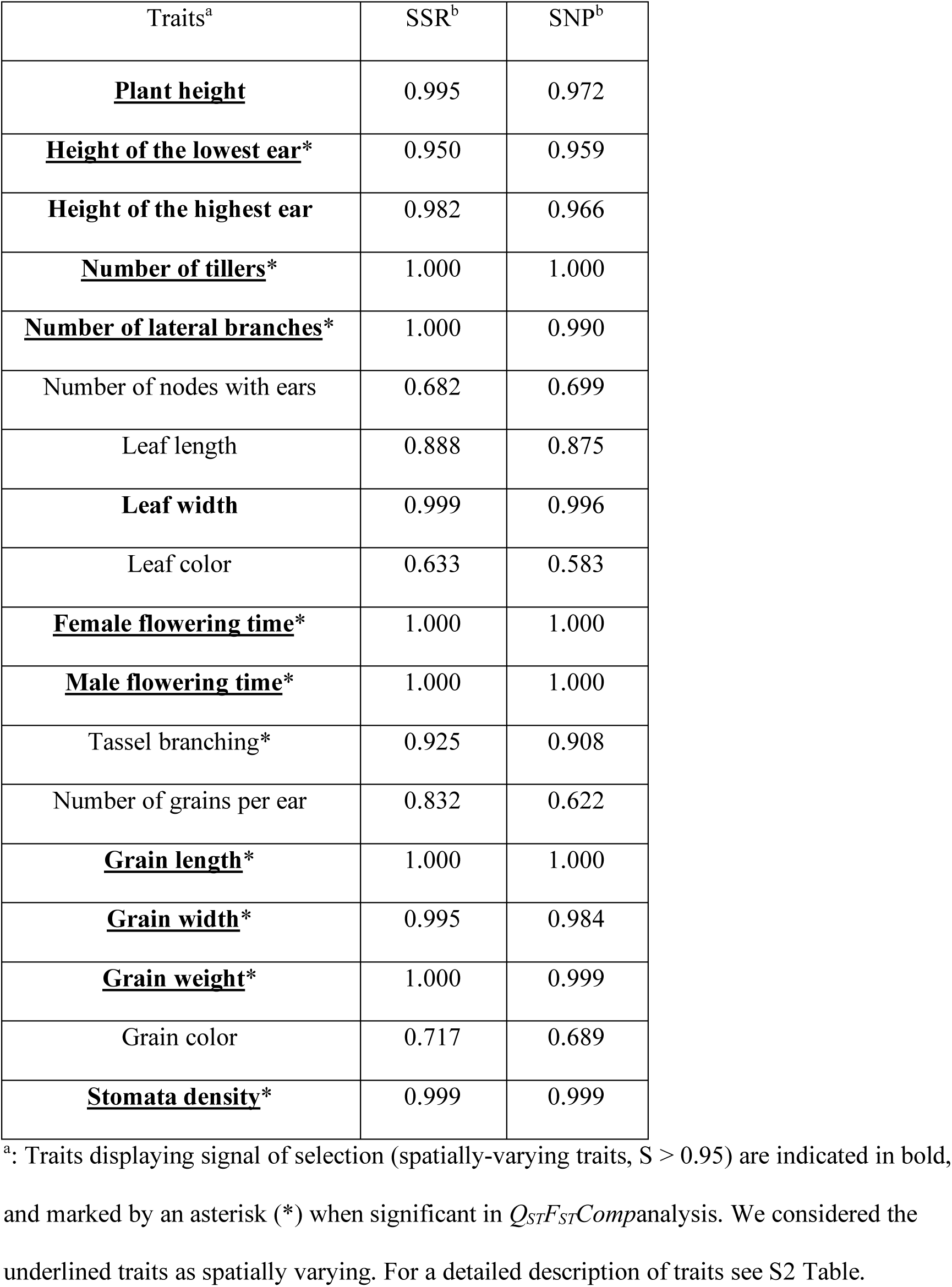
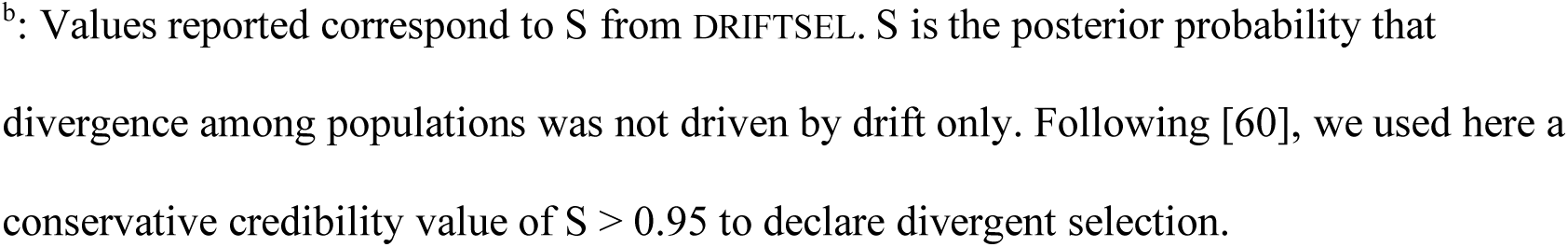
Signals of selection (posterior probability S) for each trait considering SSR markers (11 populations) or SNPs (10 populations).

| Traits<sup>a</sup> | SSR<sup>b</sup> | SNP<sup>b</sup> |
| --- | --- | --- |
| **<u>Plant height</u>** | 0.995 | 0.972 |
| **<u>Height of the lowest ear\*</u>** | 0.950 | 0.959 |
| **Height of the highest ear** | 0.982 | 0.966 |
| **<u>Number of tillers\*</u>** | 1.000 | 1.000 |
| **<u>Number of lateral branches\*</u>** | 1.000 | 0.990 |
| Number of nodes with ears | 0.682 | 0.699 |
| Leaf length | 0.888 | 0.875 |
| **Leaf width** | 0.999 | 0.996 |
| Leaf color | 0.633 | 0.583 |
| **<u>Female flowering time\*</u>** | 1.000 | 1.000 |
| **<u>Male flowering time\*</u>** | 1.000 | 1.000 |
| Tassel branching* | 0.925 | 0.908 |
| Number of grains per ear | 0.832 | 0.622 |
| **<u>Grain length\*</u>** | 1.000 | 1.000 |
| **<u>Grain width\*</u>** | 0.995 | 0.984 |
| **<u>Grain weight\*</u>** | 1.000 | 0.999 |
| Grain color | 0.717 | 0.689 |
| **<u>Stomata density\*</u>** | 0.999 | 0.999 |

<sup>a</sup>: Traits displaying signal of selection (spatially-varying traits, $S > 0.95$ ) are indicated in bold, and marked by an asterisk (\*) when significant in $Q_{ST}F_{ST}$ Comanalysis. We considered the underlined traits as spatially varying. For a detailed description of traits see S2 Table.
<sup>b</sup>: Values reported correspond to $S$ from DRIFTSEL. $S$ is the posterior probability that divergence among populations was not driven by drift only. Following [60], we used here a conservative credibility value of $S > 0.95$ to declare divergent selection.

Altogether, evidence of spatially varying selection at 10 traits (Table 1) as well as continuous variation of a subset of traits across populations in both elevation gradients (Fig 2, S3 Fig) was consistent with a syndrome where populations produced less tillers, flowered earlier, displayed lower stomata density and carried larger, longer and heavier grains with increasing elevation.

### Outlier detection and correlation with environmental variables

We successfully genotyped 218 (∼81%) out of 270 outlier SNPs on a broad set of 28 populations, of which 141 were previously detected in candidate regions for local adaptation [58].Candidate regions were originally identified from re-sequencing data of only six teosinte populations (S1 Table) following an approach that included high differentiation between highlands and lowlands, environmental correlation, and in some cases their intersection with genomic regions involved in quantitative trait variation in maize. The remaining outlier SNPs (77) were discovered in the present study by performing *F_ST_*-scans on the same re-sequencing data (S5 Table). We selected outlier SNPs that were both highly differentiated between highland and lowland populations within gradients (high/low in gradient *a* or *b* or both), and between highland and lowland populations within subspecies in gradient *b* (high/low within *parviglumis*, *mexicana* or both). *F_ST_*-scans pinpointed three genomic regions of particularly high differentiation (S9 Fig) that corresponded to previously described inversions [55, 56]: one inversion on chromosome 1 (*Inv1n*), one on chromosome 4 (*Inv4m*) and one on the far end of chromosome 9 (*Inv9e*).

A substantial proportion of outlier SNPs was chosen based on their significant correlation among six populations between variation of allele frequency and their coordinate on the first environmental principal component [58]. We extended environmental analyses to 171outlier SNPs (MAF>5%) on a broader sample of 28 populations (S1 Table) and used the two first components (PCenv1 and PCenv2) to summarize environmental information. When considering all 37 populations, the first component that explained 56% of the variation, correlated with altitude but displayed no correlation to either latitude or longitude. PCenv1 was defined both by temperature- and precipitation-related variables (S2 B Fig) including Minimum Temperature of Coldest Month (T6), Mean Temperature of Driest and Coldest Quarter (T9 and T11) and Precipitation of Driest Month and Quarter (P14 and P17). The second PC explained 20.5% of the variation and was mainly defined (S2 B Fig) by Isothermality (T3), Temperature Seasonality (T4) and Temperature Annual Range (T7).

We first employed multiple regression to test for each SNP, whether the pairwise *F_ST_*matrix across 28 populations correlated to the environmental (distance along PCenv1) and/or the geographical distance. As expected, we found a significantly greater proportion of environmentally-correlated SNPs among outliers compared with neutral SNPs (χ² =264.07, P-value=2.2 10^-16^), a pattern not seen with geographically-correlated SNPs. That outlier SNPs displayed a greater isolation-by-environment than isolation-by-distance, indicated that patterns of allele frequency differentiation among populations were primarily driven by adaptive processes. We further tested correlations between allele frequencies and environmental variation.

Roughly 60.8% (104) of the 171 outlier SNPs associated with at least one of the two first PCenvs, with 87 and 33 associated with PCenv1 and PCenv2, respectively, and little overlap (S5 Table). As expected, the principal component driven by altitude (PCenv1) correlated to allele frequency for a greater fraction of SNPs than the second orthogonal component. Interestingly, we found enrichment of environmentally-associated SNPs within inversions both for PCenv1 (χ² = 14.63, P-value=1.30 10^-4^) and PCenv2 (χ² = 33.77, P-value=6.22 10^-9^).

### Associating genotypic variation to phenotypic variation

We tested the association between phenotypes and 171 of the outlier SNPs (MAF>5%) using the association panel. For each SNP-trait combination, the sample size ranged from 264 to 1068, with a median of 1004 individuals (S6 Table). We used SSRs to correct for both structure (at *K*=5) and kinship among individual genotypes. This model (model 6) resulted in a uniform distribution of P-values when testing the association between genotypic variation at SSRs and phenotypic trait variation (S10 Fig). Under this model, we found that 126 outlier SNPs (73.7%) associated to at least one trait (Fig 5 and S11 Fig) at an FDR of 10%.The number of associated SNPs per trait varied from 0 for leaf and grain coloration, to 55 SNPs for grain length, with an average of 22.6 SNPs per trait (S5 Table). Ninety-three (73.8%) out of the 126 associated SNPs were common to at least two traits, and the remaining 33 SNPs were associated to a single trait (S5 Table). The ten traits displaying evidence of spatially varying selection in the *Q_ST_*-*F_ST_* analyses displayed more associated SNPs per trait (30.5 on average), than the non-spatially varying traits (12.75 on average).

**Figure 5:**
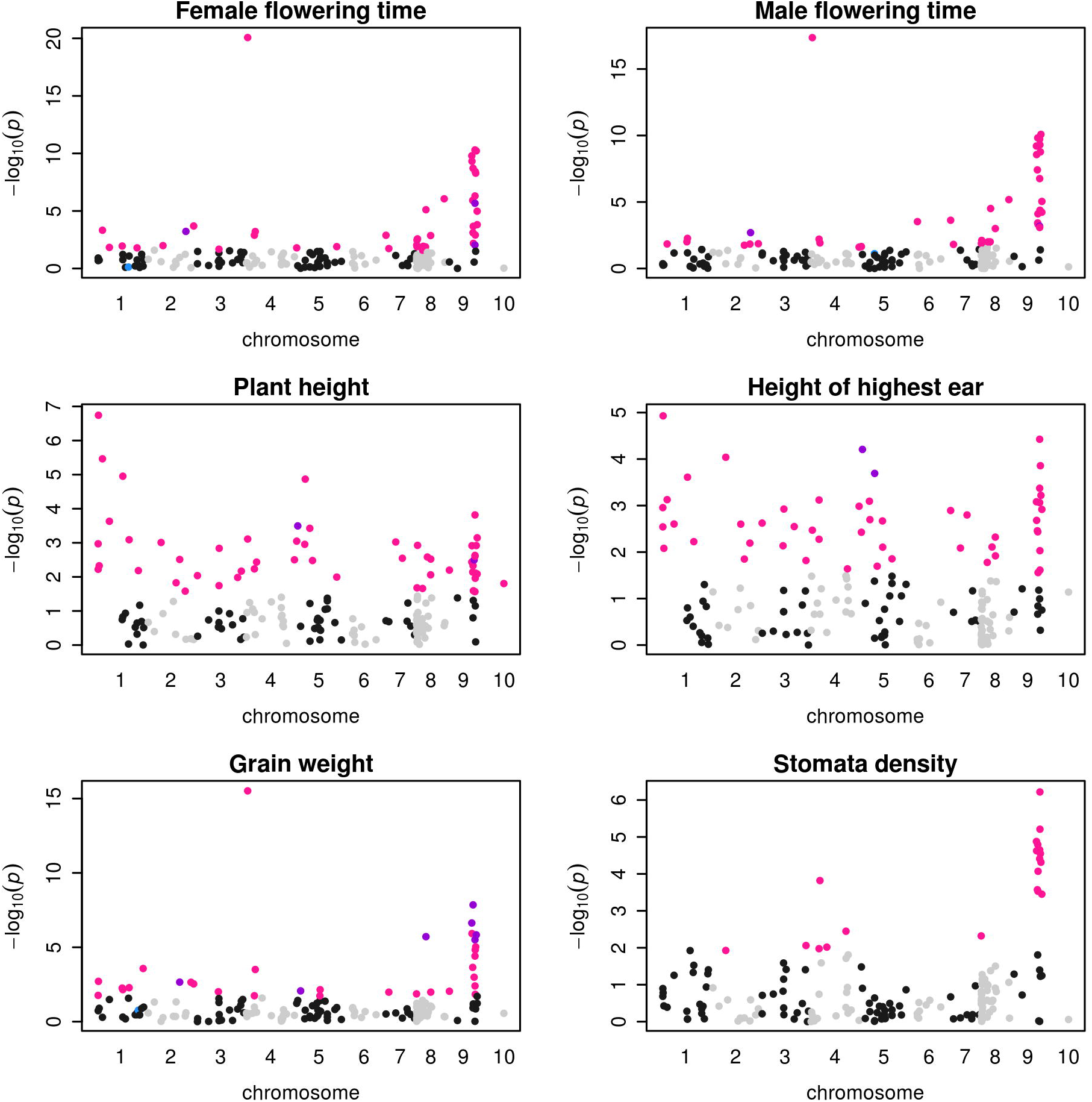
Manhattan plots of associations between 171 outlier SNPs and 6 phenotypic traits. X-axis indicates the positions of outlier SNPs on chromosomes 1 to 10, black and gray colors alternating per chromosome. Plotted on the Y-axis are the negative Log10-transformed *P* values obtained for the *K*=5 model. Significant associations (10% FDR) are indicated considering either a structure matrix at *K*=5 (pink dots), 11 populations (blue dots) or both *K*=5 and 11 populations (purple dots) models.

A growing body of literature stresses that incomplete control of population stratification may lead to spurious associations [61]. Hence, highly differentiated traits along environmental gradients are expected to co-vary with any variant whose allele frequency is differentiated along the same gradients, without underlying causal link. We therefore expected false positives in our setting where both phenotypic traits and outlier SNPs varied with altitude. We indeed found a slightly significant correlation (r=0.5, P-value=0.03) between the strength of the population effect for each trait – a measure of trait differentiation (S3 Table) – and its number of associated SNPs (S5 Table).

To verify that additional layers of structuring among populations did not cause an excess of associations, we repeated the association analyzes considering a structuring with 11 populations (instead of *K*=5) as covariate (model 7), a proxy of the structuring revealed at *K*=11 (S6 Fig). With this level of structuring, we retrieved much less associated SNPs (S5 Table). Among the 126 SNPs associating with at least one trait at *K*=5, only 22 were recovered considering 11 populations. An additional SNP was detected with structuring at 11 populations that was absent at *K*=5. Eight traits displayed no association, and the remaining traits varied from a single associated SNP (Leaf length – LeL and the number of tillers – Til) to 8 associated SNPs for grain weight (S5 Table). For instance, traits such as female or male flowering time that displayed 45 and 43 associated SNPs at *K*=5, now displayed only 4 and 3 associated SNPs, respectively (Fig 5). Note that one trait (Leaf color) associated with 4 SNPs considering 11 populations while displaying no association at *K*=5. Significant genetic associations were therefore highly contingent on the population structure. Noteworthy, traits under spatially varying selection still associated with more SNPs (2.00 on average) than those with no spatially varying selection (1.25 SNPs on average).

Altogether the 23 SNPs recovered considering a neutral genetic structure with 11 populations corresponded to 30 associations, 7 of the SNPs being associated to more than one trait (S5 Table). For all these 30 associations except in two cases (FFT with SNP_7, and MFT with SNP_28), the SNP effect did not vary among populations (non-significant SNP-by-population interaction in model 7 when we included the SNP interactions with year*field and population). For a subset of two SNPs, we illustrated the regression between the trait value and the shift of allele frequencies with altitude (Fig 6 A&B). We estimated corresponding additive and dominance effects (S7 Table). In some cases, the intra-population effect corroborated the inter-population variation with relatively large additive effects of the same sign (Fig 6). Note that in both examples shown in Fig 6, one or the other allele was dominant. In other cases, the results were more difficult to interpret with negligible additive effect but extremely strong dominance (S7 Table, SNP_210 for instance).

**Figure 6:**
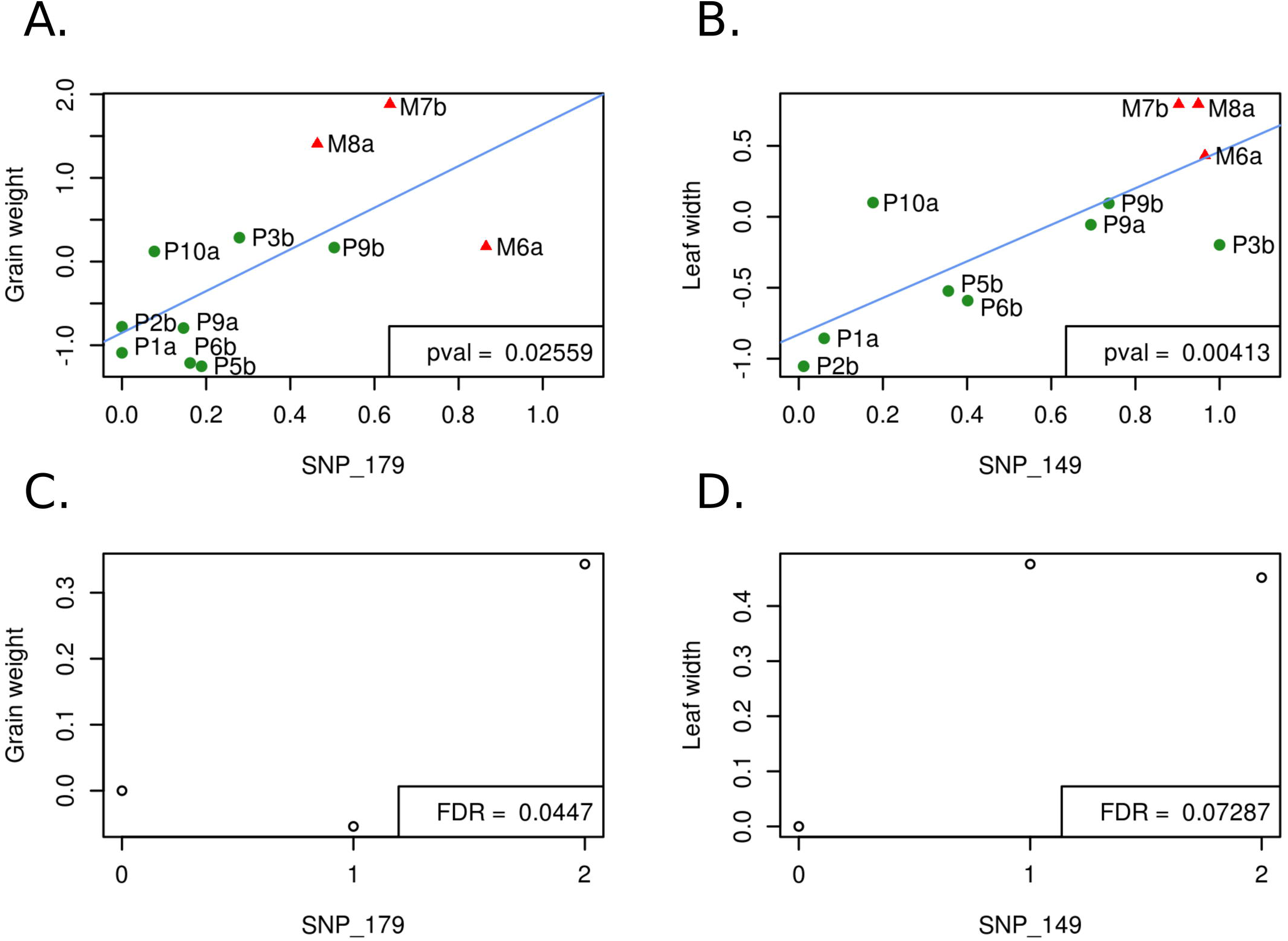
Regression of phenotypic average value on SNP allele frequency across populations, and within-populationaverage phenotypic value for each SNP genotype. Per-population phenotypic average values of traits are regressed on allele frequencies at SNP_149 (A) and SNP_179 (B) with corresponding within-population average phenotypic value per genotype (C & D). In A and B, the 11 populations of the association panel are shown with *parviglumis* (green circles) and *mexicana* (red triangles) populations sampled along gradient *a* and gradient *b*. Phenotypic average values were corrected for the experimental design (calculated as the residues of model 3). Pval refers to the P-value of the linear regression represented in blue. In C and D, genotypic effects from model 7 are expressed as the average phenotypic value of heterozygotes (1) and homozygotes for the alternative allele (2) as compared to the homozygous for the reference allele (0). FDR values were obtained from the association analysis on 171 SNPs with correction for genetic structure using 11 populations.

### Independence of SNPs associated to phenotypes

We computed the pairwise linkage disequilibrium (LD) as measured by r^2^ between the 171 outlier SNPs using the R package LDcorSV [62]. Because we were specifically interested by LD pattern between phenotypically-associated SNPs, as for the association analyses we accounted for structure (*K*=5) and kinship computed from SSRs while estimating LD [63]. The 171 outlier SNPs were distributed along the 10 chromosomes of maize, and exhibited low level of linkage disequilibrium (LD), except for SNPs located on chromosomes eight, nine, and a cluster of SNPs located on chromosome 4 (S12 Fig).

Among the 171, the subset of 23 phenotypically-associated SNPs (detected when considering the 11-populationstructure) displayed an excess of elevated LD values – out of 47 pairs of SNPs phenotypically-associated to a same trait, 16 pairs were contained in the 5% higher values of the LD distribution of all outlier SNP pairs. Twelve out of the 16 pairs associated to grain weight, the remaining four to leaf coloration, and one pair of SNPs associated to both traits. Noteworthy was that inversions on chromosomes 1, 4, and 9, taken together, were enriched for phenotypically-associated SNPs (χ² = 8.95, P-value=0.0028). We recovered a borderline significant enrichment with the correction *K*=5 (χ² = 3.82, P-value=0.051).

Finally, we asked whether multiple SNPs contributed independently to the phenotypic variation of a single trait. We tested a multiple SNP model where SNPs were added incrementally when significantly associated (FDR < 0.10). We found 2, 3 and 2 SNPs for female, male flowering time and height of the highest ear, respectively (S5 Table). For the two former traits, the SNPs were located on different chromosomes. For the latter trait, the SNPs were both located on chromosome 5 but displayed no LD (SNP_25 and SNP_30, S12 Fig).

## Discussion

Plants are excellent systems to study local adaptation. First, owing to their sessile nature, local adaptation of plant populations is pervasive [13]. Second, environmental effects can be efficiently controlled in common garden experiments, facilitating the identification of the physiological, morphological and phenological traits influenced by spatially-variable selection [64]. Identification of the determinants of complex trait variation and their covariation in natural populations is however challenging [65]. While population genomics has brought a flurry of tools to detect footprints of local adaptation, their reliability remains questioned [61]. In addition, local adaptation and demographic history frequently follow the same geographic route, making the disentangling of trait, molecular, and environmental variation, particularly arduous. Here we investigated those links on a well-established outcrossing system, the closest wild relatives of maize, along altitudinal gradients that display considerable environmental shifts over short geographical scales.

### The syndrome of altitudinal adaptation results from selection at multiple co-adapted traits

Common garden studies along elevation gradients have been conducted in European and North American plants species [66]. Together with other studies, they have revealed that adaptive responses to altitude are multifarious [67]. They include physiological responses such as high photosynthetic rates [68], tolerance to frost [69], biosynthesis of UV-induced phenolic components [70]; morphological responses with reduced stature [71, 72], modification of leaf surface [73], increase in leaf non-glandular trichomes [74], modification of stomata density; and phenological responses with variation in flowering time [75], and reduced growth period [76].

Our multivariate analysis of teosinte phenotypic variation revealed a marked differentiation between teosinte subspecies along an axis of variation (21.26% of the total variation) that also discriminated populations by altitude (Fig 2A & B). The combined effects of assortative mating and environmental elevation variation may generate, in certain conditions, trait differentiation along gradients without underlying divergent selection [77]. While we did not measure flowering time differences among populations *in situ*, we did find evidence for long distance gene flow between gradients and subspecies (Fig 4 A & C). In addition, several lines of arguments suggest that the observed clinal patterns result from selection at independent traits and is not solely driven by differences in flowering time among populations. First, two distinct methods accounting for shared population history concur with signals of spatially-varying selection at ten out of the 18 traits (Table 1). Nine of them exhibited a clinal trend of increase/decrease of population phenotypic values with elevation (S3 Fig) within at least one of the two subspecies. This number is actually conservative, because these approaches disregard the impact of selective constraints which in fact tend to decrease inter-population differences in phenotypes. Second, while male and female flowering times were positively correlated, they displayed only subtle correlations (|r|<0.16) with other spatially-varying traits except for grain weight and length (|r| <0.33). Third, we observed convergence at multiple phenotypes between the lowland populations from the two gradients that occurred despite their geographic and genetic distance (Fig 4) again arguing that local adaptation drives the underlying patterns.

Spatially-varying traits that displayed altitudinal trends, collectively defined a teosinte altitudinal syndrome of adaptation characterized by early-flowering, production of few tillers albeit numerous lateral branches, production of heavy, long and large grains, and decrease in stomata density. We also observed increased leaf pigmentation with elevation, although with a less significant signal (S3 Table), consistent with the pronounced difference in sheath color reported between *parviglumis* and *mexicana* [78, 79]. Because seeds were collected from wild populations, a potential limitation of our experimental setting is the confusion between genetic and environmental maternal effects. Environmental maternal effects could bias upward our heritability estimates. However, our results corroborate previous findings of reduced number of tillers and increased grain weight in *mexicana* compared with *parviglumis* [80]. Thus, although maternal effects could not be fully discarded, we believe they were likely to be weak.

The trend towards depleted stomata density at high altitudes (Fig 2) could arguably represent a physiological adaptation as stomata influence components of plant fitness through their control of transpiration and photosynthetic rate [81].

Indeed, in natural accessions of *A. thaliana*, stomatal traits showed signatures of local adaptation and were associated with both climatic conditions and water-use efficiency [82]. Furthermore, previous work has shown that in arid and hot highland environments, densely-packed stomata may promote increased leaf cooling in response to desiccation [83] and may also counteract limited photosynthetic rate with decreasing pCO_2_ [84]. Accordingly, increased stomata density at high elevation sites has been reported in alpine species such as the European beech [85] as well as in populations of *Mimulus guttatus* subjected to higher precipitations in the Sierra Nevada [86]. In our case, higher elevations display both *arid* environment and *cooler* temperatures during the growing season, features perhaps more comparable to other tropical mountains for which a diversity of patterns in stomatal density variation with altitude has been reported [87]. Further work will be needed to decipher the mechanisms driving the pattern of declining stomata density with altitude in teosintes. Altogether, the altitudinal syndrome was consistent with natural selection for rapid life-cycle shift, with early-flowering in the shorter growing season of the highlands and production of larger propagules than in the lowlands. This altitudinal syndrome evolved in spite of detectable gene flow.

Although we did not formally measure biomass production, the lower number of tillers and higher amount and size of grains in the highlands when compared with the lowlands may reflect trade-offs between allocation to grain production and vegetative growth [88]. Because grains fell at maturity and a single teosinte individual produces hundreds of ears, we were unable to provide a proxy for total grain production. The existence of fitness-related trade-offs therefore still needs to be formally addressed.

Beyond trade-offs, our results more generally question the extent of correlations between traits. In maize, for instance, we know that female and male flowering time are positively correlated and that their genetic control is in part determined by a common set of genes [89]. They themselves further increase with yield-related traits [90]. Response to selection for late-flowering also led to a correlated increase in leaf number in cultivated maize [91], and common genetic loci have been shown to determine these traits as well [92]. Here we found strong positive correlations between traits: male and female flowering time, grain length and width, plant height and height of the lowest or highest ear. Strong negative correlations were observed instead between grain weight and both male and female flowering time. Trait correlations were therefore partly consistent with previous observations in maize, suggesting that they were inherited from wild ancestors [93].

### Footprints of past adaptation are relevant to detect variants involved in present phenotypic variation

The overall level of differentiation in our outcrossing system (*F_ST_* ≍22%) fell close to the range of previous estimates (23% [94] and 33% [55] for samples encompassing both teosinte subspecies). It is relatively low compared to other systems such as the selfer *Arabidopsis thaliana*, where association panels typically display maximum values of *F_ST_* around 60% within 10kb-windows genome-wide [95]. Nevertheless, correction for sample structure is key for statistical associations between genotypes and phenotypes along environmental gradients. This is because outliers that display lowland/highland differentiation co-vary with environmental factors, which themselves may affect traits [96]. Consistently, we found that 73.7% SNPs associated with phenotypic variation at *K*=5, but only 13.5% of them did so when considering a genetic structure with 11 populations. Except for one, the latter set of SNPs represented a subset of the former. Because teosinte subspecies differentiation was fully accounted for at *K*=5 (as shown by the clear distinction between *mexicana* populations and the rest of the samples, Fig 4A), the inflation of significant associations at *K*=5 is not due to subspecies differentiation, but rather to residual stratification among populations within genetic groups. Likewise, recent studies in humans, where global differentiation is comparatively low [97] have shown that incomplete control for population structure within European samples strongly impacts association results [61, 98]. Controlling for such structure may be even more critical in domesticated plants, where genetic structure is inferred *a posteriori* from genetic data (rather than *a priori* from population information) and pedigrees are often not well described. Below, we show that considering more than one correction using minor peaks delivered by the Evanno statistic (S5 Fig) can be informative.

Considering a structure with 5 genetic groups, the number of SNPs associated per trait varied from 1 to 55, with no association for leaf and grain coloration (S5 Table). False positives likely represent a greater proportion of associations at *K*=5 as illustrated by a slight excess of small P-values when compared with a correction with 11 populations for most traits (S11 Fig). Nevertheless, our analysis recovered credible candidate adaptive loci that were no longer associated when a finer-grained population structure was included in the model. For instance, at *K*=5 we detected *Sugary1* (*Su1*), a gene encoding a starch debranching enzyme that was selected during maize domestication and subsequent breeding [99, 100]. We found that *Su1* was associated with variation at six traits (male and female flowering time, tassel branching, height of the highest ear, grain weight and stomata density) pointing to high pleiotropy. A previous study reported association of this gene to oil content in teosintes [101]. In maize, this gene has a demonstrated role in kernel phenotypic differences between maize genetic groups [102]. *Su1* is therefore most probably a true-positive. That this gene was no longer recovered with the 11-population structure correction indicated that divergent selection acted among populations. Indeed, allelic frequency was highly contrasted among populations, with most populations fixed for one or the other allele, and a single population with intermediate allelic frequency. With the 11-population correction, very low power is thus left to detect the effect of *Su1* on phenotypes.

Although the confounding population structure likely influenced the genetic associations, experimental evidence indicates that an appreciable proportion of the variants recovered with both *K*=5 and 11 populations are true-positives (S5 Table). One SNP associated with female and male flowering time, as well as with plant height and grain length (at *K*=5 only for the two latter traits) maps within the *phytochrome B2* (SNP_210; *phyB2*) gene. Phytochromes are involved in perceiving light signals and are essential for growth and development in plants. The maize gene *phyB2* regulates the photoperiod-dependent floral transition, with mutants producing early flowering phenotypes and reduced plant height [103]. Genes from the phosphatidylethanolamine-binding proteins (PEBPs) family – *Zea mays CENTRORADIALIS* (*ZCN*) family in maize – are also well-known to act as promotor and repressor of the floral transition in plants [104]. Z*CN8* is the main floral activator of maize [105], and both *ZCN8* and *ZCN5* strongly associate with flowering time variation [102, 106]. Consistently, we found associations of male and female flowering time with *PEBP18* (SNP_15). It is interesting to note that SNPs at two flowering time genes, *phyB2* and *PEBP18*, influenced independently as well as in combination both female and male flowering time variation (S5 Table).

The proportion of genic SNPs associated to phenotypic variation was not significantly higher than that of non-genic SNPs (i.e, SNPs >1kb from a gene) (χ²(df=1) = 0.043, P-value = 0.84 at *K*=5 and χ²(df=1)=1.623, P-value =0.020 with 11 populations) stressing the importance of considering both types of variants [107]. For instance, we discovered a non-genic SNP (SNP_149) that displayed a strong association with leaf width variation as well as a pattern of allele frequency shift with altitude among populations (Fig 6B).

### Physically-linked and independent SNPs both contribute to the establishment of adaptive genetic correlations

We found limited LD among our outlier SNPs (S12 Fig) corroborating previous reports (LD decay within <100bp, [58, 94]). However, the subset of phenotypically-associated SNPs displayed greater LD, a pattern likely exacerbated by three Mb-scale inversions located on chromosomes 1 (*Inv1n*), 4 (*Inv4m*) and 9 (*Inv9e*) that, taken together, were enriched for SNPs associated with environmental variables related to altitude and/or SNPs associated with phenotypic variation. Previous work [55, 56] has shown that *Inv1n* and *Inv4m* segregate within both *parviglumis* and *mexicana*, while two inversions on chromosome 9, *Inv9d* and *Inv9e*, are present only in some of the highest *mexicana* populations; such that all four inversions also follow an altitudinal pattern. Our findings confirmed that three of these inversions possessed an excess of SNPs with high *F_ST_* between subspecies and between low- and high-*mexicana* populations for *Inv9e* [57]. Noteworthy *Inv9d* contains a large ear leaf width quantitative trait locus in maize [107]. Corroborating these results, we found consistent association between the only SNP located within this inversion and leaf width variation in teosinte populations (S5 Table). Overall, our results further strengthen the role of chromosomal inversions in teosinte altitudinal adaptation.

Because inversions suppress recombination between inverted and non-inverted genotypes, their spread has likely contributed to the emergence and maintenance of locally adaptive allelic combinations in the face of gene flow, as reported in a growing number of other models (reviewed in [108]) including insects [109], fish [110], birds [111] and plants [26, 112]. But we also found three cases of multi-SNP determinism of traits (male and female flowering time and height of the highest ear, Table S5) supporting selection on genetically independent loci. Consistently with Weber et al. [101], we found that individual SNPs account for small proportions of the phenotypic variance (S7 Table). Altogether, these observations are consistent with joint selection of complex traits determined by several alleles of small effects, some of which being maintained in linkage through selection of chromosomal rearrangements.

## Conclusion

Elevation gradients provide an exceptional opportunity for investigating variation of functional traits in response to continuous environmental factors at short geographical scales. Here we documented patterns indicating that local adaptation, likely facilitated by the existence of chromosomal inversions, allows teosintes to cope with specific environmental conditions in spite of gene flow. We detected an altitudinal syndrome in teosintes composed of sets of independent traits evolving under spatially-varying selection. Because traits co-varied with environmental differences along gradients, however, statistical associations between genotypes and phenotypes largely depended on control of population stratification. Yet, several of the variants we uncovered seem to underlie adaptive trait variation in teosintes.

Adaptive teosinte trait variation may be relevant for maize evolution and breeding. Whether the underlying SNPs detected in teosintes bear similar effects in maize or whether their effects differ in domesticated backgrounds will have to be further investigated.

## Material and Methods

### Ethics Statement

All the field work has been done in Mexico in collaboration with Instituto Nacional de InvestigacionesForestales, Agrícolas y Pecuarias, Celaya in Celaya.

### Description of teosinte populations and sampling

We used 37 teosinte populations of *mexicana* (16) and *parviglumis* (21) subspecies from two previous collections [57, 58, 113] to design our sampling. These populations (S1 Table) are distributed along two altitudinal gradients (Fig 1). We plotted their altitudinal profiles using R ‘raster’ package [114] (S1 Fig). We further obtained 19 environmental variable layers from http://idrisi.uaemex.mx/distribucion/superficies-climaticas-para-mexico. These high-resolution layers comprised monthly values from 1910 to 2009 estimated via interpolation methods [115]. We extracted values of the 19 climatic variables for each population (S1 Table). Note that high throughput sequencing (HTS) data were obtained in a previous study for six populations out of the 37 (M6a, P1a, M7b, P2b, M1b and P8b; Fig 1, S1 Table) to detect candidate genomic regions for local adaptation [58]. The four highest and lowest of these populations were included in the association panel described below.

We defined an association panel of 11 populations on which to perform a genotype-phenotype association study (S1 Table). Our choice was guided by grain availability as well as the coverage of the whole climatic and altitudinal ranges.

Hence, we computed Principal Component Analyses (PCA) from environmental variables using the FactoMineR package in R [116] and added altitude to the PCA graphs as a supplementary variable. Our association panel comprised five populations from a first gradient (*a*) – two *mexicana* and three *parviglumis,* and six populations from a second gradient (*b*) – one *mexicana* and five *parviglumis* (Fig 1).

Finally, we extracted available SNP genotypes generated with the MaizeSNP50 Genotyping BeadChipfor 28 populations out of our 37 populations [57] (S1 Table). From this available SNP dataset, we randomly sampled 1000 SNPs found to display no selection footprint [57], hereafter neutral SNPs. Data for neutral SNPs (Data S1) are available at: https://doi.org/10.6084/m9.figshare.9901472. We used this panel of 28 populations to investigate correlation with environmental variation. Note that 10 out of the 28 populations were common to our association panel, and genotypes were available for 24 to 34 individuals per population, albeit different from the ones of our association mapping panel.

### Common garden experiments

We used two common gardens for phenotypic evaluation of the association panel (11 populations). Common gardens were located at INIFAP (Instituto Nacional de Investigaciones Forestales, Agrícolas y Pecuarias) experimental field stations in the state of Guanajuato in Mexico, one in Celaya municipality at the Campo Experimental Bajío (CEBAJ) (20°31’20’’ N, 100°48’44’’W) at 1750 meters of elevation, and one in San Luis de la Paz municipality at the Sitio Experimental Norte de Guanajuato (SENGUA) (21°17’55’’N, 100°30’59’’W) at 2017 meters of elevation. These locations were selected because they present intermediate altitudes (S1 Fig). The two common gardens were replicated in 2013 and 2014.

The original sampling contained 15 to 22 mother plants per population. Eight to 12 grains per mother plant were sown each year in individual pots. After one month, seedlings were transplanted to the field. Each of the four fields (2 locations, 2 years) was divided into four blocks encompassing 10 rows and 20 columns. We evaluated one offspring of ∼15 mother plants from each of the 11 teosinte populations in each block, using a semi-randomized design, i.e. each row containing one or two individuals from each population, and individuals being randomized within row, leading to a total of 2,640 individual teosinte plants evaluated.

### SSR genotyping and genetic structuring analyses on the association panel

In order to quantify the population structure and individual kinship in our association panel, we genotyped 46 SSRs (microsatellites, S4 Table). Primers sequences are available from the maize database project [117] and genotyping protocol were previously published [118]. Genotyping was done at the GENTYANE platform (UMR INRA 1095, Clermont-Ferrand, France). Allele calling was performed on electropherograms with GeneMapper Software Applied Biosystems. Allele binning was carried out using Autobin software [119], and further checked manually.

We employed STRUCTURE Bayesian classification software to compute a genetic structure matrix on individual genotypes. Individuals with over 40% missing data were excluded from analysis. We applied the same criterion on SSRs success rate and restricted all analyses to a subset of 38 SSRs (S4 Table). For each number of clusters (*K* from 2 to 13), we performed 10 independent runs of 500,000 iterations after a burn-in period of 50,000 iterations, and combined these 10 replicates using the LargeKGreedy algorithm from the CLUMPP program [120]. We plotted the resulting clusters using DISTRUCT software. We then used the Evanno method [121] to choose the optimal *K* value.

We inferred a kinship matrix **K** from the same SSRs using SPAGeDI [122]. Kinship coefficients were calculated for each pair of individuals as the correlation between allelic states [123]. Since teosintes are outcrossers and expected to exhibit an elevated level of heterozygosity, we estimated intra-individual kinship to fill in the diagonal. We calculated ten kinship matrices, each excluding the SSRs from one out of the 10 chromosomes. Microsatellite data (Data S2) are available at: https://doi.org/10.6084/m9.figshare.9901472.

In order to gain insights into population history of divergence and admixture, we used 1000 neutral SNPs (i.e. SNPs genotyped by Aguirre-Liguori and collaborators [57] and that displayed patterns consistent with neutrality among 49 teosinte populations) genotyped on 10 out of the 11 populations of the association panel to run a TreeMix analysis (TreeMix version 1.13 [124]). TreeMix models genetic drift to infer populations’ splits from an outgroup as well as migration edges along a bifurcating tree. We oriented the SNPs using the previously published MaizeSNP50 Genotyping BeadChip data from the outgroup species *Tripsacumdactyloides* [55]. We tested from 0 to 10 migration edges. We fitted both a simple exponential and a non-linear least square model (threshold of 1%) to select the optimal number of migration edges as implemented in the OptM R package [125]. We further verified that the proportion of variance did not substantially increase beyond the optimal selected value.

### Phenotypic trait measurements

We evaluated a total of 18 phenotypic traits on the association panel (S2 Table). We measured six traits related to plant architecture (PL: Plant Height, HLE: Height of the Lowest Ear, HHE: Height of the Highest Ear, Til: number of Tillers, LBr: number of Lateral Branches, NoE: number of Nodes with Ears), three traits related to leave morphologies (LeL: Leaf Length, LeW: Leaf Width, LeC: Leaf Color), three traits related to reproduction (MFT: Male Flowering Time, FFT: Female Flowering Time, TBr : Tassel Branching), five traits related to grains (Gr: number of Grains per ear, GrL: Grain Length, GrWi: Grain Width, GrWe: Grain Weight, GrC: Grain Color), and one trait related to Stomata (StD: Stomata Density). These traits were chosen because we suspected they could contribute to differences among teosinte populations based on a previous report of morphological characterization of 112 teosintes grown in five localities [126].

We measured the traits related to plant architecture and leaves after silk emergence. Grain traits were measured at maturity. Leaf and grain coloration were evaluated on a qualitative scale. For stomata density, we sampled three leaves per plant and conserved them in humid paper in plastic bags. Analyses were undertaken at the Institute for Evolution and Biodiversity (University of Münster) as follows: 5mm blade discs were cut out from the mid length of one of the leaves and microscopic images were taken after excitation with a 488nm laser. Nine locations (0.15mm^2^) per disc were captured with 10 images per location along the z-axis (vertically along the tissue). We automatically filtered images based on quality and estimated leaf stomata density using custom image analysis algorithms implemented in Matlab. For each sample, we calculated the median stomata density over the (up to) nine locations. To verify detection accuracy, manual counts were undertaken for 54 random samples.

Automatic and manual counts were highly correlated (R²=0.82), indicating reliable detection (see S1 Annex Stomata Detection, Dittberner and de Meaux, for a detailed description). The filtered data set of phenotypic measurements (Data S3) is available at: https://doi.org/10.6084/m9.figshare.9901472.

### Statistical analyses of phenotypic variation

In order to test for genetic effects on teosinte phenotypic variation, we decomposed phenotypic values of each trait considering a fixed population effect plus a random mother-plant effect (model 1):

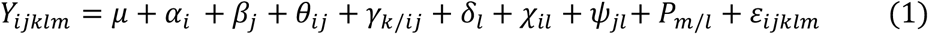

where the response variable Y is the observed phenotypic value, µ is the total mean, α*i* is the fixed year effect (*i* = 2013, 2014), β*j* the fixed field effect (*j* = field station: *SENGUA, CEBAJ)*, θ*ij* is the year-by-field interaction, γ*k/ij* is the fixed block effect (*k* = 1, 2, 3, 4) nested within the year-by-field combination, δ*l* is the fixed effect of the population of origin (*l* = 1 to 11),χ*il* is the year-by-population interaction, ψ*jl* is the field-by-population interaction, P*m/l* is the random effect of mother plant (*m* = 1 to 15) nested within population, and *εijklm* is the individual residue. Identical notations were used in all following models. For the distribution of the effects, the same variance was estimated within all populations. Mixed models were run using ASRemlv.3.0 [127] and MM4LMM v2.0.1 [https://rdrr.io/cran/MM4LMM/man/MM4LMM-package.html, update by F. Laporte] R packages, which both gave very similar results, and fixed effects were tested through Wald tests.

For each trait, we represented variation among populations using box-plots on mean values per mother plant adjusted for the experimental design following model 2:

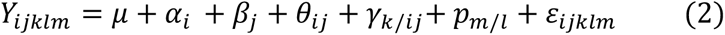

where mother plant within population is considered as a fixed effect. We used the function *predict* to obtain least-square means (ls-means) of each mother plant, and looked at the tendencies between population’s values. All fixed models were computed using *lm* package in R, and we visually checked the assumptions of residues independence and normal distribution.

We performed a principal component analysis (PCA) on phenotypic values corrected for the experimental design, using FactoMineR package in R [116] from the residues of model 3 computed using the lm package in R:

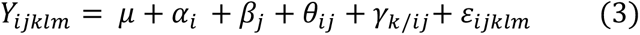

Finally, we tested for altitudinal effects on traits by considering the altitude of the sampled population (*l*) as a covariate (ALT) and its interaction with year and field in model 4:

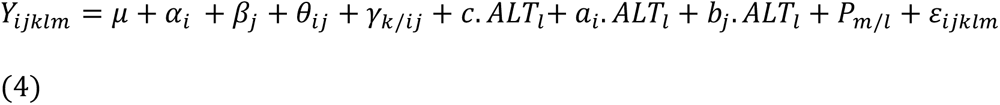

where all terms are equal to those in model 1 except that the fixed effect of the population of origin was replaced by a regression on the population altitude (ALT*l*).

### Detection of selection acting on phenotypic traits

We aimed at detecting traits evolving under spatially varying selection by comparing phenotypic to neutral genotypic differentiation. *Q_ST_* is a statistic analogous to *F_ST_*but for quantitative traits, which can be described as the proportion of phenotypic variation explained by differences among populations [19, 128].

Significant differences between *Q_ST_*and *F_ST_* can be interpreted as evidence for spatially-varying selection when *Q_ST_*>*F_ST_* [128]. We used the R package *Q_ST_F_ST_Comp*[129] that is adequate for experimental designs with randomized half-sibs in outcrossing species. We used individuals that were both genotyped and phenotyped on the association panel to establish the distribution of the difference between statistics (*Q_ST_*-*F_ST_*) under the neutral hypothesis of evolution by drift - using the half-sib dam breeding design and 1000 resamples. We next compared it to the observed difference with 95% threshold cutoff value in order to detect traits under spatially-varying selection.

In addition to *Q_ST_*-*F_ST_*analyses, we employed the DRIFTSEL R package [130] to test for signal of selection of traits while accounting for drift-driven population divergence and genetic relatedness among individuals (half-sib design). DRIFTSEL is a Bayesian method that compares the probability distribution of predicted and observed mean additive genetic values. It provides the S statistic as output, which measures the posterior probability that the observed population divergence arose under divergent selection (S∼1), stabilizing selection (S∼0) or genetic drift (intermediate S values) [59]. It is particularly powerful for small datasets, and can distinguish between drift and selection even when *Q_ST_*-*F_ST_*are equal [59]. We first applied RAFM to estimate the *F_ST_* value across populations, and the population-by-population coancestry coefficient matrix. We next fitted both the RAFM and DRIFTSEL models with 15,000 MCMC iterations, discarded the first 5,000 iterations as transient, and thinned the remaining by 10 to provide 1000 samples from the posterior distribution. Note that DRIFTSEL was slightly modified because we had information only about the dams, but not the sires, of the phenotyped individuals. We thus modified DRIFTSEL with the conservative assumption of all sires being unrelated. Because DRIFTSEL does not require that the same individuals were both genotyped and phenotyped, we used SSRs and phenotype data of the association panel as well as the set of neutral SNPs and phenotype data on 10 out of the 11 populations. For the SNP analyses, we selected out of the 1000 neutral SNPs the 465 most informative SNPs based on the following criteria: frequency of the less common variant at least 10%, and proportion of missing data at most 1%. Finally, we estimated from DRIFTSEL the posterior probability of the ancestral population mean for each trait as well as deviations of each population from these values.

Both *Q_ST_*-*F_ST_*and DRIFTSEL rely on the assumption that the observed phenotypic variation was determined by additive genotypic variation. We thus estimated narrow-sense heritability for each trait in each population to estimate the proportion of additive variance in performance. We calculated per population narrow-sense heritabilites as the ratio of the estimated additive genetic variance over the total phenotypic variance on our common garden measurements using the MCMCglmm R package [131] where half sib family is the single random factor, and the design (block nested within year and field) is corrected as fixed factor. For three grain-related traits, we also ran the same model but including mother plants phenotypic values calculated from the remaining grains not sown. We ran 100,000 iterations with 10,000 burn-in, inverse gamma (0.001; 0.001) as priors. We then calculated the mean and standard deviation of the 11 per population *h²* estimates.

### Pairwise correlations between traits

We evaluated pairwise-correlations between traits by correlating the residues obtained from model 5, that corrects the experiment design (year, field and blocks) as well as the underlying genetic structure estimated from SSRs:

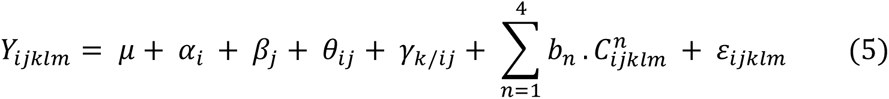

where *b_n_*is the slope of the regression of Y on the n^th^ structure covariate C*^n^*. Structure covariate values (C*^n^* covariates, from STRUCTURE output) were calculated at the individual level, i.e. for each offspring of mother plant *m* from population *l,* grown in the year *i* field *j* and block *k*. C*^n^* are thus declared with *ijklm* indices, although they are purely genetic covariates.

### Genotyping of outlier SNPs on 28 populations

We extracted total DNA from each individual plant of the association panel as well as 20 individuals from each of the 18 remaining populations that were not included in the association panel (Table 1). Extractions were performed from 30 mg of lyophilized adult leaf material following recommendations of DNeasy 96 Plant Kit manufacturer (QIAGEN, Valencia, CA, USA). We genotyped outlier SNPs using Kompetitive Allele Specific PCR technology (KASPar, LGC Group) [132]. Data for outlier SNPs (Data S4 and Data S5) are available at: https://doi.org/10.6084/m9.figshare.9901472.

Among SNPs identified as potentially involved in local adaptation, 270 were designed for KASPar assays, among which 218 delivered accurate quality data. Of the 218 SNPs, 141 were detected as outliers in two previous studies using a combination of statistical methods – including *F_ST_*-scans [133], Bayescan [32] and Bayenv2 [35, 134], Bayescenv [135] – applied to either six of our teosinte populations [58] or to a broader set of 49 populations genotyped by the Illumina MaizeSNP50 BeadChip [57]. The remaining outlier SNPs (77) were detected by *F_ST_*-scans from six populations (S9 Fig, S5 Table), following a simplified version of the rationale in [58] by considering only differentiation statistics: SNPs were selected if they displayed both a high differentiation (5% highest *F_ST_*values) between highland and lowland populations in at least one of the two gradients, and a high differentiation (5% highest *F_ST_* values) between highland and lowland populations either within *parviglumis* (P2b and P8b) or within *mexicana* (M7b and M1b) or both in gradient *b* (Fig 1). We thereby avoided SNPs fixed between the two subspecies.

### Association mapping

We tested the association of phenotypic measurements with outlier SNPs on a subset of individuals for which (1) phenotypic measurements were available, (2) at least 60% of outlier SNPs were adequately genotyped, and (3) kinship and cluster membership values were available from SSR genotyping. For association, we removed SNPs with minor allele frequency lower than 5%.

In order to detect statistical associations between outlier SNPs and phenotypic variation, we used the following mixed model derived from [136]:

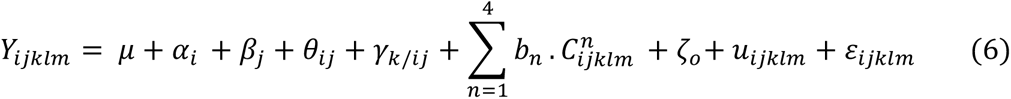

where ζ is the fixed bi-allelic SNP factor with one level for each of the three genotypes (*o*=0, 1, 2; with *o*=1 for heterozygous individuals), and u*ijklm* is the random genetic effect of the individual. We assumed that the vector of u*ijklm* effects followed a *N*(0,**K** σ^2^u) distribution, where **K** is the kinship matrix computed as described above.

A variant of model 6 was employed to test for SNP association to traits, while correcting for structure as the effect of population membership (*δl*), *δ* being a factor with 11 levels (populations):

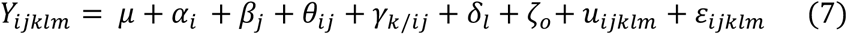

In order to avoid overcorrection of neutral genetic structure and improve power, we ran the two models independently for each chromosome using a kinship matrix **K** estimated from all SSRs except those contained in the chromosome of the tested SNP [137]. We tested SNP effects through the Wald statistics, and applied a 10% False Discovery Rate (FDR) threshold for each phenotype separately. In order to validate the correction for genetic structure, the 38 multiallelic SSR genotypes were transformed into biallelic genotypes, filtered for MAF > 5%, and used to run associations with the complete 6 and 7 models, as well as 6-type models excluding either kinship or both structure and kinship. For each trait, we generated QQplots of P-values for each of these models.

Multiple SNP models were built by successively adding at each step the most significant SNP, as long as its FDR was lower than 0.10. We controlled for population structure considering 11 populations and used the kinship matrix that excluded the SSRs on the same chromosome as the last tested SNP.

### Environmental correlation of outlier SNPs

We tested associations between allelic frequency at 171 outlier SNPs and environmental variables across 28 populations, using Bayenv 2.0 [40]. Because environmental variables are highly correlated, we used the first two principal component axes from the environmental PCA analysis (PCenv1 and PCenv2) to run Bayenv 2.0. This software requires a neutral covariance matrix, that we computed from the available dataset of 1000 neutral SNPs (S1 Table). We performed 100,000 iterations, saving the matrix every 500 iterations. We then tested the correlation of these to the last matrix obtained, as well as to an *F_ST_* matrix calculated with BEDASSLE [138], as described in [57].

For each outlier SNP, we compared the posterior probability of a model that included an environmental factor (PCenv1 or PCenv2) to a null model. We determined a 5% threshold for significance of environmental association by running 100,000 iterations on neutral SNPs. We carried out five independent runs for each outlier SNP and evaluated their consistency from the coefficient of variation of the Bayes factors calculated among runs.

In order to test whether environmental distance was a better predictor of allele frequencies at candidate SNPs than geography, we used multiple regression on distance matrices (MRM, [139]) implemented in the ecodist R package [140] for each outlier SNP. We used pairwise *F_ST_*values as the response distance matrix and the geographic and environmental distance matrices as explanatory matrices. We evaluated the significance of regression coefficients by 1000 permutations and iterations of the MRM. We determined the total number of environmentally and geographically associated SNPs (P-value<0.05) among outliers. We employed the same methodology for our set of 1000 neutral SNPs.

## Supporting information

Annex 1

## Acknowledgments

We thank Jessica Melique for her help in gathering stomata data. We are very grateful to Angelica Cibrian at Langebio (Cinvestav, Irapuato, Mexico) who let us use her lab to lyophilize our samples, and store them. Valeria Souza greatly helped us with logistic support in Mexico. Insights from Delphine Legrand, Pierre de Villemereuil and Elodie Marchadier were important for analyzing correlations between traits and to estimate heritabilities. We would like to thank warmly five anonymous reviewers for their insightful comments on our work. GQE-Le Moulon benefits from the support of Saclay Plant Sciences-SPS (ANR-17-EUR-0007).

## Supporting information captions

**Figure S1: Altitudinal profiles along gradients *a* and *b*.** Sampled populations are plotted on parallel altitudinal profiles for gradients *a* and *b*. Darker gray lines indicate lower latitude for gradient *a* and lower longitude for gradient *b*. Sampled populations are plotted by green circles (*parviglumis*) or red triangles (*mexicana*). The altitude of the two experimental fields (CEBAJ: 1750m and SENGUA: 2017m) are marked with asterisks on the y-axes.

**Figure S2: Principal Component Analysis of 19 climate variables for 37 teosinte populations.** A: Projection of *parviglumis* (in green) and *mexicana* (in red) populations on the first PCA plane with gradients *a* and *b* indicated by triangles and circles, respectively. The 11 populations evaluated in common gardens are surrounded by a purple outline. Populations that were previously sequenced to detect selection footprints are shown in bold (S1 Table). B: Correlation circle of the 19 climatic variables on the first PCA plane. Climatic variables indicated as Tn (n from 1 to 11) and Pn (n from 12 to 19) are related to temperature and precipitation, respectively. Altitude, Latitude and Longitude (in blue) were added as supplementary variables, and CEBAJ and SENGUA field locations were added as supplementary individuals.

**Figure S3: Box-plots of means adjusted by field, year and block, for all traits.** Populations are ranked by altitude. *parviglumis* populations are shown in green and *mexicana* in red. Lighter colors are used for gradient ‘a’ and darker colors for gradient ‘b’. Units of measurement correspond to those defined in S2 Table. For male and female flowering time, we report values for all 11 populations although very few individuals from the two most lowland populations (P1a and P2b) flowered. Covariation with altitude was significant for all traits except for the number of nodes with ears on the main tiller (S3 Table).

**Figure S4: Pairwise correlations between phenotypic traits.** Pearson coefficient sign and magnitude for significant correlations between phenotypic traits after correction for experiment design (Model 2). X: correlations that are not significant.

**Figure S5.** Evanno method calculations for population number ΔK in the association panel genotyped for 38 SSRs.

**Figure S6. Genetic clustering of ancestry proportions in the association panel genotyped for 38 SSRs.** Genetic clustering was computed for *K*=2 to *K*=11. Vertical lines (individuals) are partitioned into coloured segments whose length represents the admixture proportions from the K clusters.

**Figure S7. Determination of the migration edge number in the TreeMix model.** Observed Log likelihood values are plotted against the number of migration edges tested from 0 to 10, and two models are fitted to the data (A). Both the simple exponential and the non-linear least squares delivered an optimal value of 3 for the number of migration edges (change points). The model with 3 migration edges explained 98.75% of the variance, a substantial increase from the null model with no migration edge which is 95.7% (B).

**Figure S8: Significance of *Q_ST_*-*F_ST_* difference for each trait.** The dotted blue line indicates the 95% threshold of the simulated distributions and the red line refers to the observed difference. In this analysis, we considered as spatially-varying traits those for which the observed difference fell outside the 95% threshold. Note that Plant height was borderline significant. *: Set of traits detected by DRIFTSEL.

**Figure S9: Genomic *F_ST_*-scans on 6 teosinte populations.** We computed 4 pairwise-*F_ST_* values from 6 populations previously sequenced (S1 Table). Those include *F_ST_*between lowland and highland populations of each gradient (P1a-M6a, P2b-M7b) as well as within subspecies on gradient *b* (P2b-P8b, M1b-M7b). *F_ST_* values are averaged across sliding windows of 20 SNPs with a step of five SNPs (from top to bottom, chromosome 1 to 10) and normalized by subtracting the *F_ST_* mean and dividing by the standard deviation across pairwise comparisons. Only the top 1% values are represented. The 1‰ thresholds for each pairwise comparisons are indicated by colored horizontal lines. Horizontal black bars indicate location of inversions on chromosome 1 (*Inv1n*), chromosome 4 (*Inv4m*) and chromosome 9 (*Inv9d*). The subset of 171 outlier SNPs analyzed in the present study is indicated with black diamond marks along the X axes.

**Figure S10: QQ-plots of observed P-values and expected P-values generated from 38 SSRs.** We employed three versions of the model 6 with correction for neither structure nor kinship, with correction for genetic structure (at *K*=5), with correction for genetic structure (at *K*=5 and with 11 populations) and kinship.

**Figure S11: Manhattan plots of associations between 171 outlier SNPs and 12 phenotypic traits.** X-axis indicates the positions of outlier SNPs on chromosomes 1 to 10, black and gray colors alternating per chromosome. Plotted on the Y-axis are the negative Log_10_-transformed *P* values obtained for the *K*=5 model. Significant associations (10% FDR) are indicated considering either a structure matrix at *K*=5 (pink dots), for 11 populations (blue dots), or for both *K*=5 and 11 populations models (purple dots).

**Figure S12: Pairwise Linkage Disequilibrium (LD) between outlier SNPs**. Pairwise LD between 171 SNPs was estimated using r^2^, and corrected for structure at *K*=5 and kinship computed from 38 SSRs. Blue shaded bars show the 23 SNPs found to associate with at least one phenotype under the 11 populations structure correction.

**S1 Table. Description of 37 teosinte populations and sets of populations used in the present study by data types.**

**S2 Table. List of the 18 phenotypic traits measured and estimates of narrow-sense heritabilities (*h^2^*).**

**S3 Table. Significance of main effects for each trait as determined by models 1 and 4.**

**S4 Table. Description of 46 SSRs and genotyping success rate.**

**S5 Table. Characteristics, association with phenotypes, effects and correlation with environment of outlier SNPs.**

**S6 Table. Number of individuals used to test associations between 171 SNPs and 18 phenotypes.**

**S7 Table**. **Additive and dominance effects of SNPs associated to traits after the 11-population structure correction.**

S1 Annex. Stomata detection.

All data (Data S1-S5) are available at : https://doi.org/10.6084/m9.figshare.9901472.

Data S1. Set of neutral SNPs used for 10 populations.

Data S2. Genotyping of 38 microsatellites on the association panel.

Data S3. Phenotypic data from the association panel.

Data S4. Candidate SNPs genotyping on the association panel.

Data S5. Candidate SNPs genotyping on 28 populations.

## References

1. Whitlock MC. Modern approaches to local adaptation. The American Naturalist. 2015;186.

2. Bradshaw AD. Ecological significance of genetic variation between populations. Perspectives on plant population ecology. 1984:213–28.

3. Bulmer MGG. Multiple niche polymorphism. The American Naturalist. 1972;106:254–7.

4. Endler JA. Natural selection in the wild. 1986:354.

5. Gay L, Crochet PA, Bell DA, Lenormand T. Comparing clines on molecular and phenotypic traits in hybrid zones: A window on tension zone models. Evolution. 2008;62:2789–806.

6. Lande R. Natural selection and random genetic drift in phenotypic evolution. Evolution. 1976;30:314–34.

7. Lenormand T. Gene flow and the limits to natural selection. Trends in Ecology and Evolution. 2002;17:183–9.

8. Whitlock MC, Gomulkiewicz R. Probability of fixation in a heterogeneous environment. Genetics. 2005:1–40.

9. Yeaman S, Otto SP. Establishment and maintenance of adaptive genetic divergence under migration, selection and drift. Evolution. 2011;67:2123–9.

10. Rundle HD, Nosil P. Ecological speciation. Ecology Letters. 2005;8:336–52.

11. Kawecki TJ, Ebert D. Conceptual issues in local adaptation. Ecology Letters. 2004;7:1225–41.

12. Hereford J. A quantitative survey of local adaptation and fitness trade-offs. The American Naturalist. 2009;173:579–88.

13. Leimu R, Fischer M. A meta-analysis of local adaptation in plants. PLoS ONE. 2008;3.

14. Tiffin P, Ross-Ibarra J. Advances and limits of using population genetics to understand local adaptation. Trends in Ecology and Evolution. 2014;29:673–80.

15. Garcia-Ramos G, Kirkpatrick M. Genetic models of adaptation and gene flow in peripherial populations. Evolution. 1997;51:21–8.

16. Slatkin M. Rare alleles as indicators of gene flow. Evolution. 1985;39(1):53–65.

17. Lande R. Neutral theory of quantitative genetic variance in an island model with local extinction and colonization. Evolution. 1992;46:381–9.

18. Whitlock MC. Neutral additive genetic variance in a metapopulation. Genetics Research. 1999;74:215–21.

19. Spitze K. Population structure in *Daphina obtusa*: quantitative genetic and allozymic variation. Genetics Society of America. 1993;135:367–74.

20. Wright S. The genetical structure of populations. Annals of Eugenics. 1951;15:323–54.

21. Luquet E, Léna J-P, Miaud C, Plénet S. Phenotypic divergence of the common toad (*Bufo bufo*) along an altitudinal gradient: evidence for local adaptation. Heredity. 2015;114:69–79.

22. Roschanski AM, Csilléry K, Liepelt S, Oddou-Muratorio S, Ziegenhagen B, Huard Fdr, et al. Evidence of divergent selection for drought and cold tolerance at landscape and local scales in *Abies alba* Mill. in the French Mediterranean Alps. Molecular Ecology. 2016;25:776–94.

23. Kawakami T, Morgan TJ, Nippert JB, Ocheltree TW, Keith R, Dhakal P, et al. Natural selection drives clinal life history patterns in the perennial sunflower species, *Helianthus maximiliani*. Molecular Ecology. 2011;20:2318–28.

24. Moyers BT, Rieseberg LH. Remarkable life history polymorphism may be evolving under divergent selection in the silverleaf sunflower. Molecular Ecology. 2016;25:3817–30.

25. Kirkpatrick M, Barton N. Chromosome inversions, local adaptation and speciation. Genetics. 2006.

26. Lowry DB, Willis JH. A widespread chromosomal inversion polymorphism contributes to a major life-history transition, local adaptation, and reproductive isolation. PLoS Biology. 2010;8.

27. Legrand D, Larranaga N, Bertrand R, Ducatez S, Calvez O, Stevens VM, et al. Evolution of a butterfly dispersal syndrome. Proceedings of the Royal Society B. 2016; 283(1839): 20161533

28. Bierne N, Welch J, Loire E, Bonhomme F, David P. The coupling hypothesis: Why genome scans may fail to map local adaptation genes. Molecular Ecology. 2011;20:2044–72.

29. Lewontin RC, Krakauer J. Distribution of gene frequency as a test of the theory of the selective neutrality of polymorphisms. Genetics. 1973;74:175–95.

30. Beaumont MA, Nichols RA. Evaluating loci for use in the genetic analysis of population structure. Proceedings of the Royal Society B. 1996;263:1619–26.

31. Vitalis R, Dawson K, Boursot P. Interpretation of variation across marker loci as evidence of selection. Genetics. 2001;158:1811–23.

32. Foll M, Gaggiotti O. A genome scan method to identify selected loci appropriate for both dominant and codominant markers: A Bayesian perspective. Genetics. 2008;180:977–93.

33. Excoffier LaH, T and Foll, Matthieu. Detecting loci under selection in a hierarchically structured population. Heredity. 2009;103:285–98.

34. Bonhomme M, Chevalet C, Servin B, Boitard S, Abdallah J, Blott S, et al. Detecting selection in population trees: The Lewontin and Krakauer test extended. Genetics. 2010;186:241–62.

35. Günther T, Coop G. Robust identification of local adaptation from allele frequencies. Genetics. 2013;195:205–20.

36. Lotterhos KE, Whitlock MC. Evaluation of demographic history and neutral parameterization on the performance of *F_ST_* outlier tests. Molecular Ecology. 2014;23:2178–92.

37. Haasl RJ, Payseur BA. Fifteen years of genomewide scans for selection: trends, lessons and unaddressed genetic sources of complication. Molecular Ecology. 2016;25:5–23.

38. Le Corre V, Kremer A. The genetic differentiation at quantitative trait loci under local adaptation. Molecular Ecology. 2012;21:1548–66.

39. Yi X, Liang Y, Huerta-Sanchez E, Jin X, Xi Ping Cuo Z, Pool JE, et al. Sequencing of fifty human exomes reveals adaptations to high altitude. Science. 2010;329:75–8.

40. Coop G, Witonsky D, Di Rienzo A, Pritchard JK. Using environmental correlations to identify loci underlying local adaptation. Genetics. 2010;185:1411–23.

41. Guillot G, Renaud S, Ledevin R, Michaux J, Claude J. A unifying model for the analysis of phenotypic, genetic, and geographic data. Systematic Biology. 2012;61:897–911.

42. Frichot E, Schoville SD, Bouchard G, François O. Testing for associations between loci and environmental gradients using latent factor mixed models. Molecular Biology and Evolution. 2013;30:1687–99.

43. Gautier M. Genome-wide scan for adaptive divergence and association with population-specific covariates. Genetics. 2015;201:1555–79.

44. Joost S, Bonin A, Bruford MW, Després L, Conord C, Erhardt G, et al. A spatial analysis method (SAM) to detect candidate loci for selection: Towards a landscape genomics approach to adaptation. Molecular Ecology. 2007;16:3955–69.

45. Poncet BN, Herrmann D, Gugerli F, Taberlet P, Holderegger R, Gielly L, et al. Tracking genes of ecological relevance using a genome scan in two independent regional population samples of *Arabis alpina*. Molecular Ecology. 2010;19:2896–907.

46. De Mita S, Thuillet AC, Gay L, Ahmadi N, Manel S, Ronfort J, et al. Detecting selection along environmental gradients: Analysis of eight methods and their effectiveness for outbreeding and selfing populations. Molecular Ecology. 2013;22:1383–99.

47. Hoban S, Kelley JL, Lotterhos KE, Antolin MF, Bradburd G, Lowry DB, et al. Finding the genomic basis of local adaptation: pitfalls, practical solutions, and future directions. The American Naturalist. 2016;188:379–97.

48. Barton N, Hermisson J, Nordborg M. Why structure matters. eLife. 2019;8.

49. Fournier-Level A, Korte A, Cooper MD, Nordborg M, Schmitt J, Wilczek AM. A map of local adaptation in Arabidopsis thaliana. Science. 2011;334:86–9.

50. Hancock AM, Brachi B, Faure N, Horton MW, Jarymowycz LB, Sperone FG, et al. Adaptation to climate across the *Arabidopsis thaliana* genome. Science. 2011;334:83–6.

51. Ross-Ibarra J, Tenaillon M, Gaut BS. Historical divergence and gene flow in the genus Zea. Genetics. 2009;181:1399–413.

52. Hufford MB, Martínez-Meyer E, Gaut BS, Eguiarte LE, Tenaillon MI. Inferences from the historical distribution of wild and domesticated maize provide ecological and evolutionary insight. PLoS ONE. 2012;7.

53. Bilinski P, Albert PS, Berg JJ, Birchler JA, Grote MN, Lorant A, et al. Parallel altitudinal clines reveal trends in adaptive evolution of genome size in *Zea mays*. PLoS Genetics. 2018;14.

54. Diez CM, Gaut BS, Meca E, Scheinvar E, Montes-Hernandez S, Eguiarte LE, et al. Genome size variation in wild and cultivated maize along altitudinal gradients. New Phytologist. 2013;199:264–76.

55. Pyhäjärvi T, Hufford MB, Mezmouk S, Ross-Ibarra J. Complex patterns of local adaptation in teosinte. Genome Biology and Evolution. 2013;5:1594–609.

56. Fang Z, Pyhäjärvi T, Weber AL, Dawe RK, Glaubitz JC, Sánchez González JdJ, et al. Megabase-scale inversion polymorphism in the wild ancestor of maize. Genetics. 2012;191:883–94.

57. Aguirre-Liguori JA, Tenaillon MI, Vázquez-Lobo A, Gaut BS, Jaramillo-Correa JP, Montes-Hernandez S, et al. Connecting genomic patterns of local adaptation and niche suitability in teosintes. Molecular Ecology. 2017;26:4226–40.

58. Fustier MA, Brandenburg JT, Boitard S, Lapeyronnie J, Eguiarte LE, Vigouroux Y, et al. Signatures of local adaptation in lowland and highland teosintes from whole-genome sequencing of pooled samples. Molecular Ecology. 2017;26:2738–56.

59. Ovaskainen O, Karhunen M, Zheng C, Arias JMC, Merilä J. A new method to uncover signatures of divergent and stabilizing selection in quantitative traits. Genetics. 2011;189:621–32.

60. McKinney GJ, Varian A, Scardina J, Nichols KM. Genetic and morphological divergence in three strains of brook trout Salvelinus fontinalis commonly stocked in Lake Superior. PLoS ONE. 2014;9(12):e113809.

61. Sohail M, Maier RM, Ganna A, Bloemendal A, Martin AR, Turchin MC, et al. Polygenic adaptation on height is overestimated due to uncorrected stratification in genome-wide association studies. eLife. 2019;8.

62. Desrousseaux AD, Sandron F, Siberchicot A, Cierco-Ayrolles C, Mangin B. R Package ‘ LDcorSV ’. 2017.

63. Mangin B, Siberchicot A, Nicolas S, Doligez A, This P, Cierco-Ayrolles C. Novel measures of linkage disequilibrium that correct the bias due to population structure and relatedness. Heredity. 2012;108:285–91.

64. Savolainen O, Lascoux M, Merilä J. Ecological genomics of local adaptation. Nature reviews Genetics. 2013;14:807–20.

65. Anderson JT, Willis JH, Mitchell-Olds T. Evolutionary genetics of plant adaptation. Trends in Genetics. 2011;27:258–66.

66. Halbritter AH, Fior S, Keller I, Billeter R, Edwards PJ, Holderegger R, et al. Trait differentiation and adaptation of plants along elevation gradients. Journal of Evolutionary Biology. 2018;31(6):784–800.

67. Körner C. The use of ‘altitude’ in ecological research. Trends in Ecology and Evolution. 2007;22:569–74.

68. Friend AD, Woodward FI, Switsur VR. Field measurements of photosynthesis, stomatal conductance, leaf nitrogen and δ 13 C along altitudinal gradients in Scotland. Functional Ecology. 1989;3:117.

69. Neuner G. Frost resistance in alpine woody plants. Frontiers in Plant Science. 2014;5.

70. Frohnmeyer H, Staiger D. Update on ultraviolet-B light responses ultraviolet-B radiation-mediated responses in plants. Balancing damage and protection. Plant Physiology. 2014;133:1420–8.

71. Byars SG, Papst W, Hoffmann AA. Local adaptation and cogradient selection in the alpine plant, *Poa hiemata*, along a narrow altitudinal gradient. Evolution. 2007;61:2925–41.

72. Luo Y, Widmer A, Karrenberg S. The roles of genetic drift and natural selection in quantitative trait divergence along an altitudinal gradient in *Arabidopsis thaliana*. Heredity. 2015;114:220–8.

73. Guerin GR, Wen H, Lowe AJ, Guerin GR, Wen H. Leaf morphology shift linked to climate change. Population Ecology. 2012:882–6.

74. Kofidis G, Bosabalidis AM, Moustakas M. Contemporary seasonal and altitudinal variations of leaf structural features in oregano (*Origanum vulgare* L.). Annals of Botany. 2003;92(5):635–45.

75. Mendez-Vigo B, Pico FX, Ramiro M, Martinez-Zapater JM, Alonso-Blanco C. Altitudinal and climatic adaptation is mediated by flowering traits and FRI, FLC, and PHYC genes in *Arabidopsis*. Plant Physiology. 2011;157:1942–55.

76. Oleksyn J, Modrzynski J, Tjoelker MG, Zytkowaik R, Reich PB, Karolewski P. Growth and physiology of *Picea abies* populations from elevational transects: common garden evidence for altitudinal ecotypes and cold adaptation. Functional Ecology. 1998:573–90.

77. Soularue JP, Kremer A. Evolutionary responses of tree phenology to the combined effects of assortative mating, gene flow and divergent selection. Heredity. 2014;113:485–94.

78. Doebley JF. Maize introgression into teosinte - a reappraisal. Annals of the Missouri Botanical Garden. 1984;71:1100–13.

79. Lauter N, Gustus C, Westerbergh A, Doebley J. The inheritance and evolution of leaf pigmentation and pubescence in teosinte. Genetics. 2004;167:1949–59.

80. Smith JSC, Goodman MM, Lester RN. Variation within teosinte. I. Numerical analysis of morphological data. Economic Botany. 1981;35:187–203.

81. Raven J, A. Selection pressures on stomatal evolution. New Phytologist. 2002;153:371–86.

82. Dittberner H, Korte A, Mettler-Altmann T, Weber APM, Monroe G, de Meaux J. Natural variation in stomata size contributes to the local adaptation of water-use efficiency in *Arabidopsis thaliana*. Molecular Ecology. 2018:4052–65.

83. Carlson JE, Adams CA, Holsinger KE. Intraspecific variation in stomatal traits, leaf traits and physiology reflects adaptation along aridity gradients in a South African shrub. Annals of Botany. 2016;117:195–207.

84. Körner C, Mayr R. Stomatal behaviour in alpine plant communities between 600 and 2600 metres above sea level. In: Grace J, Ford ED, Jarvis PG, editors. Plants and their Atmospheric Environment: Blackwell, Oxford; 1981. p. pp 205–18.

85. Bresson CC, Vitasse Y, Kremer A, Delzon S. To what extent is altitudinal variation of functional traits driven by genetic adaptation in European oak and beech? Tree Physiology. 2011;31:1164–74.

86. Kooyers NJ, Greenlee AB, Colicchio JM, Oh M, Blackman BK. Replicate altitudinal clines reveal that evolutionary flexibility underlies adaptation to drought stress in annual *Mimulus guttatus*. New Phytologist. 2015;206:152–65.

87. Körner C, Neumayer M, Menendez-Riedl SP, Smeets-Scheel A. Functional morphology of mountain plants. Flora. 1989;182:353–83.

88. Jakobsson A, Eriksson O. A comparative study of seed number, seed size, seedling size and recruitment in grassland plants. Oikos. 2000;88:494–502.

89. Buckler ES, Holland JB, Bradbury PJ, Acharya CB, Brown PJ, Browne C, et al. The genetic architecture of maize flowering time. Science. 2009;325:714–8.

90. Moreau L, Charcosset A, Gallais A. Use of trial clustering to study QTL x environment effects for grain yield and related traits in maize. Theoretical and Applied Genetics. 2004;110:92–105.

91. Durand E, Bouchet S, Bertin P, Ressayre A, Jamin P, Charcosset A, et al. Flowering time in maize: Linkage and epistasis at a major effect locus. Genetics. 2012;190:1547–62.

92. Li D, Wang X, Zhang X, Chen Q, Xu G, Xu D, et al. The genetic architecture of leaf number and its genetic relationship to flowering time in maize. New Phytologist. 2015;210:256–68.

93. Yang CJ, Samayoa LF, Bradbury PJ, Olukolu BA, Xue W, York AM, et al. The genetic architecture of teosinte catalyzed and constrained maize domestication. Proceedings of the National Academy of Sciences of the United States of America. 2019; 116(12):5643–5652.

94. Aguirre-Liguori JA, Gaut BS, Jaramillo-Correa JP, Tenaillon MI, Montes-Hernández S, García-Oliva F, et al. Divergence with gene flow is driven by local adaptation to temperature and soil phosphorus concentration in teosinte subspecies (*Zea mays parviglumis* and *Zea mays mexicana*) Molecular Ecology. 2019:2814–30.

95. Consortium G. 1,135 Genomes reveal the global pattern of polymorphism in *Arabidopsis thaliana*. Cell. 2016;166:481–91.

96. Novembre J, Barton NH. Tread lightly interpreting polygenic tests of selection. Genetics. 2018; 208(4):1351–5.

97. Guo J, Wu Y, Zhu Z, Zheng Z, Trzaskowski M, Zeng J, et al. Global genetic differentiation of complex traits shaped by natural selection in humans. Nature Communications. 2018;9(1):1865.

98. Berg JJ, Harpak A, Sinnott-Armstrong N, Joergensen AM, Mostafavi H, Field Y, et al. Reduced signal for polygenic adaptation of height in UK Biobank. eLife. 2019;8:1–47.

99. Whitt SR, Wilson LM, Tenaillon MI, Gaut BS, Buckler ES. Genetic diversity and selection in the maize starch pathway. Proceedings of the National Academy of Sciences of the United States of America. 2002;99:12959–62.

100. Jaenicke-Despres V, Buckler ES, Smith BD, Gilbert MTP, Cooper A, Doebley J, et al. Early allelic selection in maize as revealed by ancient DNA. Science. 2003;302:1206–9.

101. Weber AL, Briggs WH, Rucker J, Baltazar BM, De Jesús Sánchez-González J, Feng P, et al. The genetic architecture of complex traits in teosinte (*Zea mays* ssp. *parviglumis*): New evidence from association mapping. Genetics. 2008;180:1221–32.

102. Bouchet S, Servin B, Bertin P, Madur D, Combes V, Dumas F, et al. Adaptation of maize to temperate climates: Mid-density genome-wide association genetics and diversity patterns reveal key genomic regions, with a major contribution of the Vgt2 (ZCN8) locus. PLoS ONE. 2013;8.

103. Sheehan MJ, Kennedy LM, Costich DE, Brutnell TP. Subfunctionalization of PhyB1 and PhyB2 in the control of seedling and mature plant traits in maize. Plant Journal. 2007;49:338–53.

104. Danilevskaya ON, Meng X, Hou Z, Ananiev EV, Simmons CR. A genomic and expression compendium of the expanded PEBP gene family from maize. Plant Physiology. 2007;146:250–64.

105. Meng X, Muszynski MG, Danilevskaya ON. The FT-Like ZCN8 gene functions as a floral activator and is involved in photoperiod sensitivity in maize. The Plant Cell. 2011;23:942–60.

106. Li YX, Li C, Bradbury PJ, Liu X, Lu F, Romay CM, et al. Identification of genetic variants associated with maize flowering time using an extremely large multi-genetic background population. The Plant journal. 2016;86:391–402.

107. Yu J, Li X, Zhu C, Yeh C-T, Wu W, Takacs E, et al. Genic and non-genic contributions to natural variation of quantitative traits in maize. Genome research. 2012:2436–44.

108. Wellenreuther M, Bernatchez L. Eco-evolutionary genomics of chromosomal inversions. Trends in Ecology and Evolution. 2018;33:427–40.

109. Ayala D, Ullastres A, González J. Adaptation through chromosomal inversions in *Anopheles*. Frontiers in Genetics. 2014;5:1–10.

110. Barth JMI, Berg PR, Jonsson PR, Bonanomi S, Corell H, Hemmer-Hansen J, et al. Genome architecture enables local adaptation of Atlantic cod despite high connectivity. Molecular Ecology. 2017;26:4452–66.

111. Lundberg M, Liedvogel M, Larson K, Sigeman H, Grahn M, Wright A, et al. Genetic differences between willow warbler migratory phenotypes are few and cluster in large haplotype blocks. Evolution Letters. 2017:155–68.

112. Twyford AD, Friedman J. Adaptive divergence in the monkey flower *Mimulus guttatus* is maintained by a chromosomal inversion. Evolution. 2015;69:1476–86.

113. Díez CM, Gaut BS, Meca E, Scheinvar E, Montes-Hernandez S, Eguiarte LE, et al. Genome size variation in wild and cultivated maize along altitudinal gradients. New Phytologist. 2013;199(1):264–76.

114. Hijmans RJ, van Etten J, Cheng J, Mattiuzzi M, Sumner M, Greenberg JA, et al. Package ‘raster’: geographic data analysis and modeling. 2018.

115. Cuervo-Robayo AP, Téllez-Valdés O, Gómez-Albores MA, Venegas-Barrera CS, Manjarrez J, Martínez-Meyer E. An update of high-resolution monthly climate surfaces for Mexico. International Journal of Climatology. 2014;34(7):2427–37.

116. Husson F, Josse J, Le S, Mazet J. Package ‘ FactoMineR ’. An R package. 2016:96.

117. Andorf CM, Cannon EK, Portwood JL, Gardiner JM, Harper LC, Schaeffer ML, et al. MaizeGDB update: New tools, data and interface for the maize model organism database. Nucleic Acids Research. 2016;44:1195–201.

118. Camus-Kulandaivelu L, Veyrieras JB, Madur D, Combes V, Fourmann M, Barraud S, et al. Maize adaptation to temperate climate: Relationship between population structure and polymorphism in the Dwarf8 gene. Genetics. 2006;172:2449–63.

119. Guichoux E, Lagache S, Wagner S, Chaumeil P, Léger P, Lepais O, et al. Current trends in microsatellite genotyping. Molecular Ecology Resources. 2011;11:591–611.

120. Jakobsson M, Rosenberg NA. CLUster Matching and Permutation Program Version 1.1.2. 2007.

121. Evanno G, Regnaut S, Goudet J. Detecting the number of clusters of individuals using the software STRUCTURE: A simulation study. Molecular Ecology. 2005;14:2611–20.

122. Hardy OJ, Vekemans X. spagedi: a versatile computer program to analyse spatial genetic structure at the individual or population levels. Molecular Ecology Notes. 2002;2:618–20.

123. Loiselle BA, Sork VL, Nason JD, Graham C. Spatial genetic structure of a tropical understory shrub, *Psychotria officinalis* (Rubiaceae). American Journal of Botany. 1995;82:1420–5.

124. Pickrell JK, Pritchard JK. Inference of population splits and mixtures from genome-wide allele frequency data. PLOS Genetics. 2012;8(11):e1002967.

125. Fitak RRs. optM: an R package to optimize the number of migration edges using threshold models. Journal of Heredity. 2019.

126. Sanchez JdJ, Kato Yamakake TA, Aguilar Sanmiguel M, Hernandez Casillas JM, Lopez Rodriguez A, Ruiz Corral JA. Distribución y caracterización del teocintle. 1998:165.

127. Butler D, Cullis BR, Gilmour AR, Gogel BJ. ASReml-R reference manual. Technical Report. 2007.

128. Holsinger KE, Weir BS. Genetics in geographically structured populations: defining, estimating and interpreting *F_ST_*. Nature reviews Genetics. 2009;10:639–50.

129. Gilbert KJ, Whitlock MC. QST-FST comparisons with unbalanced half-sib designs. Molecular Ecology Resources. 2015;15:262–7.

130. Karhunen M, Merilä J, Leinonen T, Cano JM, Ovaskainen O. driftsel: An R package for detecting signals of natural selection in quantitative traits. Molecular Ecology Resources. 2013;13:746–54.

131. Hadfield JD. MCMC methods for multi-response generalized linear mixed models: the MCMCglmm R package. Journal of Statistical Software. 2010;33.

132. Semagn K, Babu R, Hearne S, Olsen M. Single nucleotide polymorphism genotyping using Kompetitive Allele Specific PCR (KASP): Overview of the technology and its application in crop improvement. Molecular Breeding. 2014;33:1–14.

133. Weir BS, Hill WG. Estimating F-statistics. Annu Rev Genet. 2002;36:721–50.

134. Günther T, Coop G. A Short Manual for Bayenv2.0. 2016.

135. De Villemereuil P, Gaggiotti OE. A new FST-based method to uncover local adaptation using environmental variables. Methods in Ecology and Evolution. 2015.

136. Yu J, Pressoir G, Briggs WH, Vroh Bi I, Yamasaki M, Doebley JF, et al. A unified mixed-model method for association mapping that accounts for multiple levels of relatedness. Nature Genetics. 2006;38(2):203–8.

137. Rincent R, Moreau L, Monod H, Kuhn E, Melchinger AE, Malvar RA, et al. Recovering power in association mapping panels with variable levels of linkage disequilibrium. Genetics. 2014;197:375–87.

138. Bradburd GS, Ralph PL, Coop GM. Disentangling the effects of geographic and ecological isolation on genetic differentiation. Evolution. 2013;67:3258–73.

139. Lichstein JW. Multiple regression on distance matrices: A multivariate spatial analysis tool. Plant Ecology. 2007;188:117–31.

140. Goslee SC, Urban DL. The ecodist package for dissimilarity-based analysis of ecological data. Journal of Statistical Software. 2007.

