## Supplementary material for "Common gardens in teosintes reveal the establishment of a syndrome of adaptation to altitude": Annex 1

### Stomata detection in Teosinte

#### Microscopic imaging

Leaf samples were stored at 4°C during the timeframe of imaging. From each sample one leaf (of three per bag) was used. From the leaf one 5mm disc was cut for microscopic analysis. The disc was cut from the middle (vertically) of the leaf. The leaf discs were put into 80-well glass bottom plates and pressed against the well bottom using a custom-made, spring-mounted stamp array. The samples were stained with Calcofluor + 10% KOH (1:1) in order to increase cell wall fluorescence. Microscopic images of the leaf discs were taken in high throughput using the Opera High Content Screening System. A 20x water-immersion objective was used. A 488nm laser was used for excitation and fluorescence was captured at 520nm. For each disc, 9 images of pre-defined locations were taken, hereinafter referred to as image "fields". Each field represents an area of 0.15mm². In each of these locations 10 images were taken in a stack along the z-Axis to counter the height variability on the leaf surface. This resulted in 9 stacks per sample and a total of 24,480 stacks.

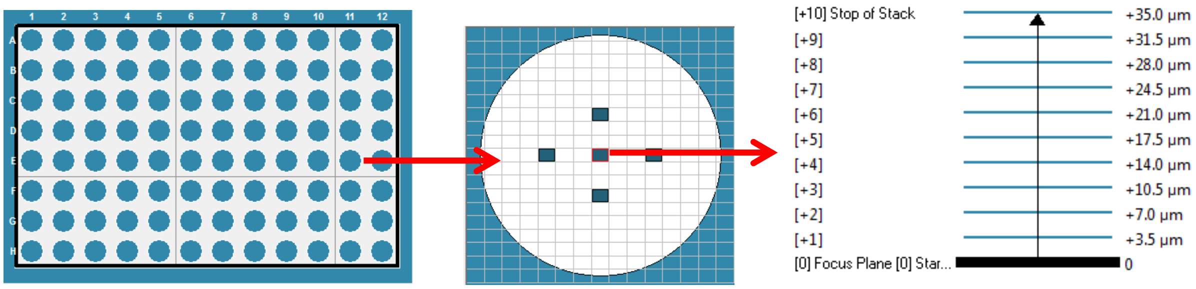


*Figure 1: Scheme of the different levels of image acquisition. First picture shows the 96-well plate (first and last column could not be used for technical reason). Second picture shows a single well with 5 image field selected for acquisition. Last picture shows the z-stack of images for the central image field.*

#### Image Analysis

The image analysis algorithms were implemented in Matlab. For stomata detection, the image "stacks" were collapsed into single 2D images by maximum intensity projection, and then saved in bitmap format. Additionally, a second composite image with enhanced cell walls was created for each stack. Therefor each layer was filtered to reduce background signal and increase contrast and was excluded from the stack if the cell wall to background ratio was too low.

#### Detection of stomata

A large fraction of images were not suited for stomata analysis due to disturbing factors caused by the nature of the samples, e.g. leaf veins, molding or surface height variation. Therefore, images were automatically selected based on the median brightness of 9 image blocks: The median brightness of at least 7 blocks had to be greater than 80. The cutoff for this was set based on two sets of manually selected good quality images and bad quality images, respectively. In high quality images stomata appeared as black holes in a white surface of epidermal cells (Figure 3, top-left). Thus, for initial object detection a simple intensity threshold was used: all pixels lower than 30 were set to one and all others set to 0, thus creating a binary image with white pixels (1) as foreground object and black pixels (0) as background (Figure 3, top-right). Then a number of filters were applied to these preliminary objects: First, very small objects (<100px) were removed. Then adjacent objects were merged by dilation followed by erosion in order to connect the two holes that form one stoma. Then objects that were too small (<300px) or too large (<3000px) for stomata were removed (Figure 3, bottom-left). The median size of the remaining preliminary stomata was calculated. In order to be considered as true stomata objects had to meet the following requirements: First, the object's area should be greater than the median area - 500px and smaller than the median area +1200px, but at least 600px. The ellipse representing the object should have a major axis shorter than 3000px and its eccentricity should be smaller than 0.92. All objects passing this filter were considered to be true stomata. As a final check of the detection quality the image was separated in 16 blocks. Only if in at least 13 of these blocks stomata were found the detection was considered successful (Figure 3, bottom-right). Otherwise the results of the image were discarded. This is due to the fact that stomata are generally distributed evenly throughout the image and if this is not the case it is likely because of out of focus areas or disturbing objects in the image. Because area of the image was known ($0.15mm^2$), the stomata counts were converted to stomatal density ($stomata/mm^2$). Then for each sample the median and standard deviation of all measured fields was calculated. In order to test the accuracy of the algorithm, stomata were counted manually for 54 random samples (median of 9 images per sample). These manual counts were then tested for correlation with the automatic measurements. The correlation coefficient $R^2$ is 0.82, indicating good correlation between the two methods.

#### Detection of cells

For detection of cells the image with emphasized cell walls was used (Figure 4, top-left). First, image quality was checked using a score for binary thresholding. If this score was smaller than 0.75 the image was discarded. Otherwise, a Canny edge detection function was applied, resulting in a binary image with edges (high contrast areas) marked as white single pixel lines (Figure 4, top-right). This means that each cell wall surrounded by a double line. In order to merge these lines the image was dilated and holes within the cell wall were removed. Then the cell wall was thinned to an equal thickness (Figure 4, bottom-left). The image was inverted so cells became foreground objects. Cells were filtered to be larger than 2500px and smaller than 10,000px. The objects that passed this filter were considered to be true cells (Figure 4, bottom-right). Due to disturbances in the cell wall intensity cells were not closed in all parts of the image and a reliable edge connection algorithm could not be developed. Therefore, cell density was estimated from only the cells that could be detected in the image. For all detected cells the total area was calculated. The number of cells was then divided by the total area to obtain an estimate of cell density in the image. In order to test the accuracy of the algorithm, cells were counted manually on 53 random samples (median of 9 images per sample). These manual counts were then tested for correlation with the automatic measurements. The correlation coefficient $R^2$ is 0.81, indicating good correlation between the two methods.

### Output

For reliable detection of stomata and cells high image quality was crucial. Due to the nature of the samples and the high throughput imaging approach this could not always be achieved. Therefore stomatal density could only be measured in 59% of the 2800 samples. In 41% of these 1670 samples cell density was successfully estimated. Results were saved as a table in csv format with the following columns: Sample ID, Median stomatal density [stomata/mm$^2$], Std. dev. of stomatal density [stomata/mm$^2$], Manual control of stomatal density [stomata/mm$^2$] Number of analyzed fields, Median cell density [cells/mm$^2$] and Manual control of cell density [cells/mm$^2$].

### Figures

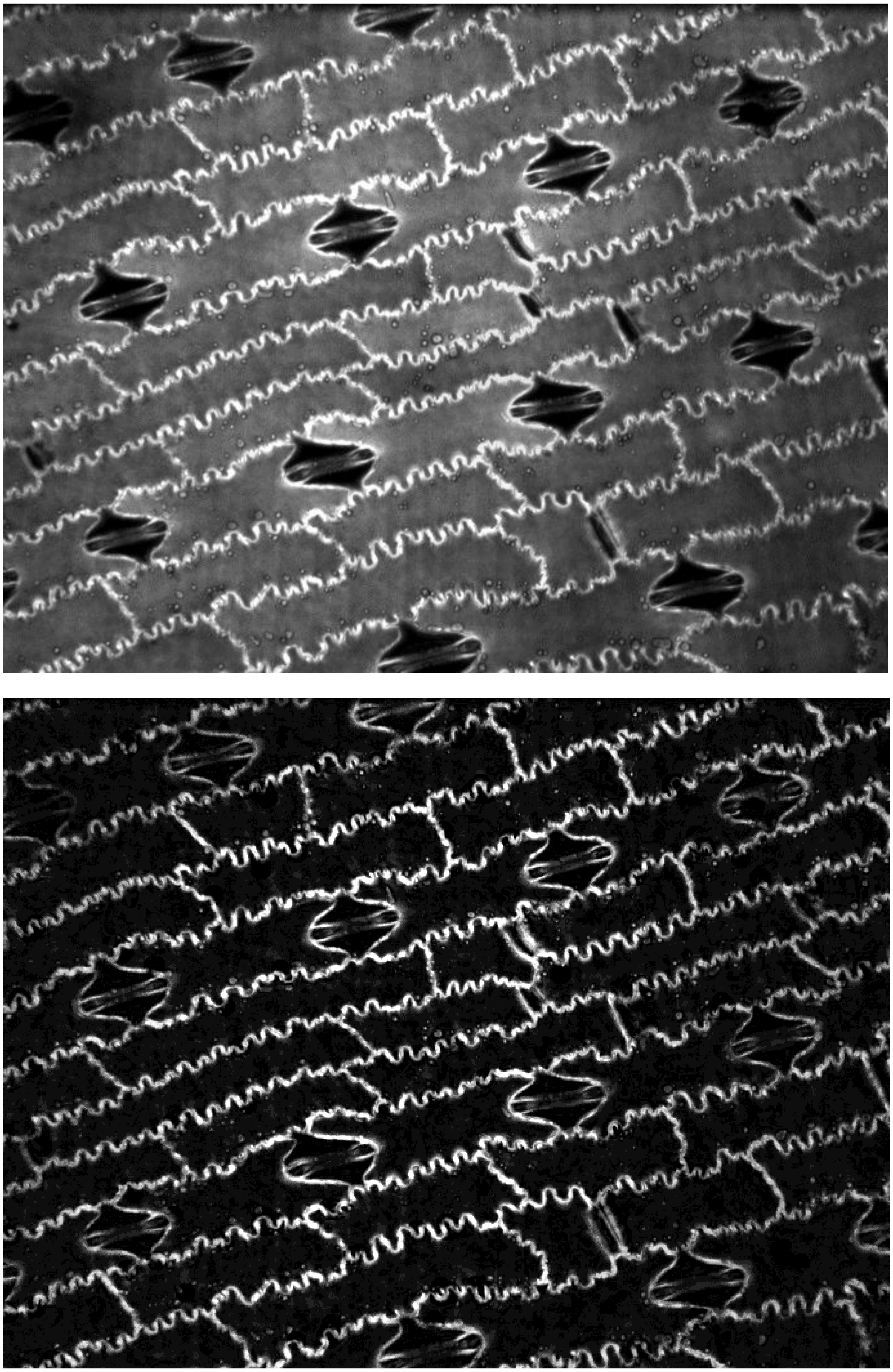

*Figure 2: Comparison between image for stomata detection (top) and image for cell detection with emphasized cell walls (bottom).*

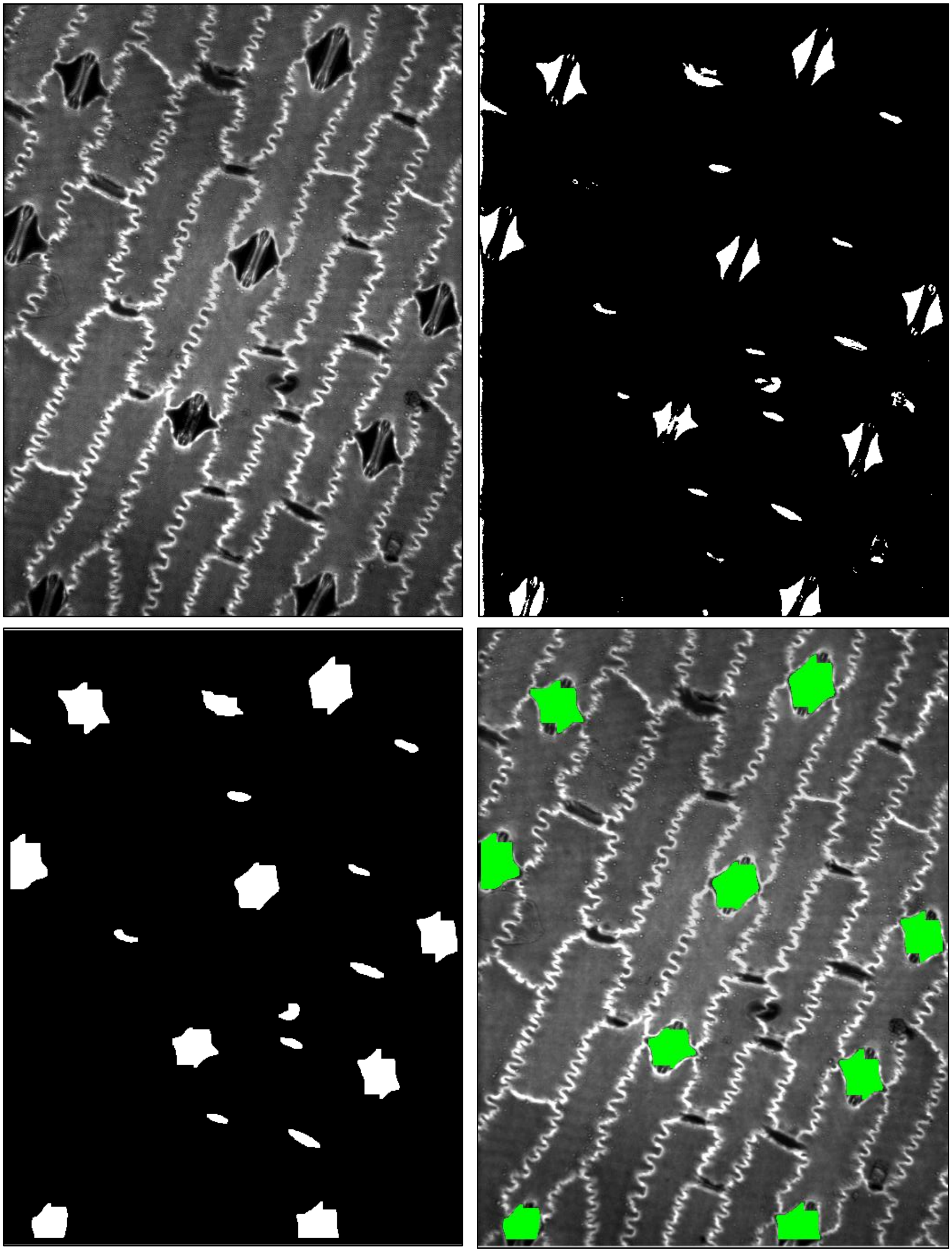

*Figure 3: Stages of stomata detection: top-left: original image; top-right: initial binary image; bottom-left: merged objects after first filter; bottom-right: overlay of image and detected stomata (green).*

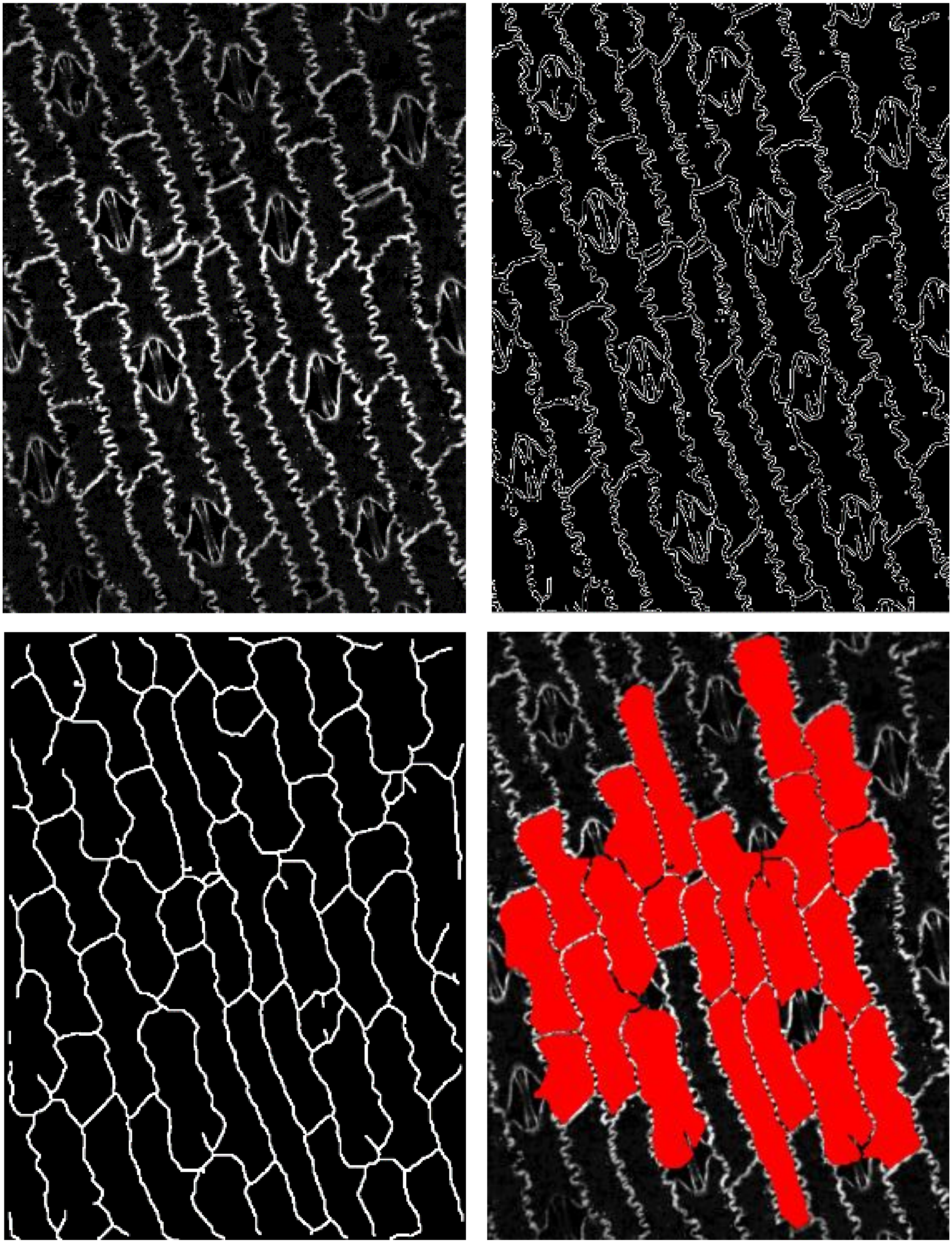

*Figure 4: Stages of cell detection: top-left: original image with emphasized cell walls; top-right: edge detection; bottom-left: detected cell walls; bottom-right: overlay of image and detected cells (red).*
